# Characterization of BLUF-photoreceptors present in *Acinetobacter nosocomialis*

**DOI:** 10.1101/2021.07.23.453528

**Authors:** Inés Abatedaga, Bárbara Pérez Mora, Marisel Tuttobene, Gabriela Müller, Daiana Biancotti, Claudio D. Borsarelli, Lorena Valle, Maria A. Mussi

**Affiliations:** Instituto de Bionanotecnología del NOA (INBIONATEC-CONICET), Universidad Nacional de Santiago del Estero (UNSE), Santiago del Estero, Argentina; Centro de Estudios Fotosintéticos y Bioquímicos (CEFOBI-CONICET), Universidad Nacional de Rosario (UNR), Rosario, Argentina

## Abstract

*Acinetobacter nosocomialis* is a Gram-negative opportunistic pathogen, whose ability to cause disease in humans is well recognized. Blue light has been shown to modulate important physiological traits related to persistence and virulence in this microorganism. In this work, we characterized the three Blue Light sensing Using FAD (BLUF) domain-containing proteins encoded in the *A. nosocomialis* genome, which account for the only “traditional” light sensors present in this microorganism. By focusing on a light-modulated bacterial process such as motility, the temperature dependence of light regulation was studied, as well as the expression pattern and spectroscopic characteristics of the different *A. nosocomialis* BLUFs. Our results show that the three BLUF-containing proteins encode active photoreceptors, despite only two of them are stable in the light-regulatory temperature range when expressed recombinantly. *In vivo*, only the *A. baumannii*’s ortholog AnBLUF65 is expressed, which is active in a temperature range from 15 °C to 37 °C. In turn, AnBLUF46 is an active photoreceptor between 15 °C to 32 °C *in vitro*, but is not expressed in *A. nosocomialis* in the conditions tested. Intra-protein interactions were analyzed using 3D models built based on *A. baumanni*’s photoreceptor, to support spectroscopic data and profile intra-protein residue interactions. A general scheme is presented on how hydrophobic/aromatic interactions may contribute to the stability of dark/light- adapted states, indicating the importance of these interactions in BLUF photoreceptors.

## Introduction

*Acinetobacter nosocomialis* is a Gram-negative coccobacillus, member of the *Acinetobacter calcoaceticus-Acinetobacter baumannii* (ACB) complex [1]. While *A. baumannii* predominates over all other members of the ACB complex in terms of incidence, poorer clinical outcomes, and antibiotic resistance rates, *A. nosocomialis* has gained recognition also as a clinically relevant human pathogen [2].

Although *Acinetobacter* spp. are primarily associated with pneumonia, they are also frequent causes of wound and burn infections, meningitis, urinary tract infections, and sepsis [3]. There is a growing trend for these isolates to display high levels of antibiotic resistance, with some being resistant to all clinically available antibiotics [4].

The ability of *A. nosocomialis* to cause disease in humans is well-recognized [5–7]. Many potential virulence factors have been identified in *A. nosocomialis* and include a CTFR inhibitory factor (Cif), a protein O-glycosylation system, a type-I secretion system, a type-II secretion system, secretion of outer membrane vesicles, the OmpA protein, the CpaA protease, and quorum sensing [2].

We have previously shown that *A. nosocomialis* is able to sense and respond to light modulating biofilm formation and motility at 24 °C [8]. Also, we have shown that light modulates persistence, metabolism, the ability to grow under iron limiting conditions and virulence in this microorganism [9]. The genome of *A. nosocomialis* RUH2624 encodes three Blue Light sensing Using FAD (BLUF)- domain containing proteins, as the only “traditional” light sensors [8]. Extensive work performed on *A. baumannii* showed that this microorganism encodes only one BLUF-type photoreceptor, designated BlsA, which functions at low-environmental temperatures up to 24 °C and is regulated both at the transcriptional level as well as the activity of the photocycle [10, 11]. Also other BLUFs from *Acinetobacter* have been characterized based on light induced phenotypes, gene knockouts and transcriptomic analyses [12, 13]. In this work, we present evidence indicating that regulation of motility by light in *A. nosocomialis* is maintained in a wide range of temperatures from 24 to 37 °C. Recombinant expression, purification and characterization of the different BLUF-domain containing-proteins showed that the three of them encode active photoreceptors; however only AnBLUF46 and AnBLUF65 are stable. Interestingly, only *anbluf65* is expressed *in vivo* and exhibits a stable photocycle in the temperature-range at which light regulates motility in *A. nosocomialis*. Spectroscopic characterization and analyses of 3D models built for these proteins provide insights into the intra-protein signaling process connecting the widely characterized BLUF photophysics with the subtle re-arrangements located in the C-terminal part of these proteins. Finally, proteomic analyses revealed that light mainly regulates proteins related to signalling and cellular metabolism.

## Material and Methods

### Bacterial Strains, Plasmids, and Media

Luria-Bertani (LB) broth (Difco) and agar (Difco) were used to grow and maintain bacterial strains. Broth cultures were incubated at the indicated temperatures either statically or with shaking at 200 rpm.

### Blue light treatments

Blue light treatments were performed as described in our previous studies [8-11, 14-18]. Briefly, cells were incubated at different temperatures in the dark or under blue light emitted by 9-light-emitting diode (LED) arrays, with an intensity of 6 to 10 μmol photons m^-2^ s^-1^. Each array was built using 3-LED module strips emitting blue light, with emission peaks centered at 462 nm, determined using the LI-COR LI-1800 spectroradiometer [15].

### Cell Motility assays

Cell motility was tested on 1% tryptone, 0.5% NaCl, 0.3% agarose plates inoculated on the surface by depositing 3 μl of Tryptic Soy Broth (TSB) cultures grown to an optical density at 660 nm (OD_660_) of 0.3. The plates were incubated in the presence or absence of blue light at the indicated temperatures for 24 hours or else specified. Three independent experiments were performed.

### Analyses of gene expression by qRT-PCR

Retrotranscription and qRT-PCR analysis were done as described in Tuttobene et al., 2019, using primers listed in Table 1. Data are presented as NRQ (Normalized Relative Quantities) calculated by the qBASE method [19], using *recA* and *rpoB* genes as normalizers. The *anbluf65* transcript levels of each sample were normalized to the *rpoB* transcript level for each cDNA sample. Relative gene expression to *rpoB* was calculated using the comparative 2^-ΔCT^ method [20]. Each cDNA sample was run in technical triplicate and repeated in at least three independent sets of samples. *t*-test was used to determine whether two values were significantly different comparing data within each temperature assayed. *p*-values: *, p < 0.01; **, p < 0.001.

**Table 1.**
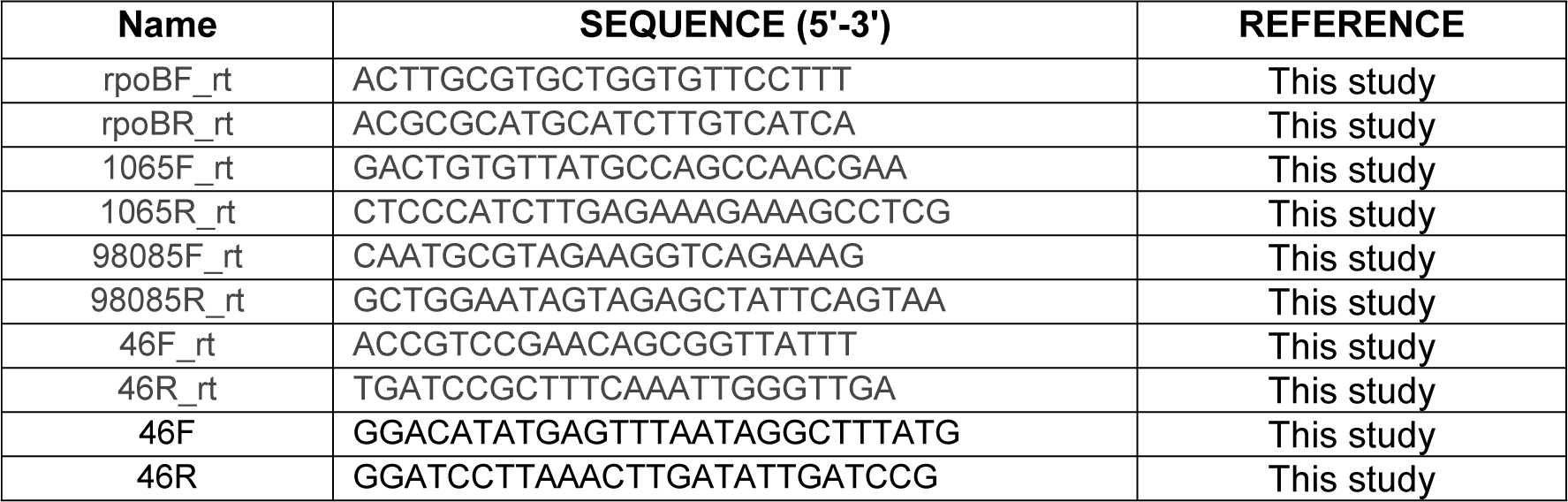
Primers used in this study.

### Cloning, synthesis, overexpression and purification of AnBLUFs

In the case of gene *anbluf46,* a PCR product amplified from *A. nosocomialis* RUH2624 genomic DNA using primers 46F and 46R (Table 1), which contain *Nde*I and *BamH*I restriction sites, respectively, was cloned into pGEM®T- Easy vector and then subcloned into the *Nde*I and *BamH*I sites of pET28-TEV. In turn, *anbluf65* and *anbluf85* coding sequences were directly synthesized and subcloned into pET 28 a(+) (Genscript, USA). Thus, the different proteins were overexpressed as N-terminal His-tag fusions. Plasmids were transformed into *E. coli* BL21 (pLys) cells, which were cultured in LB broth supplemented with chloramphenicol and kanamycin at 37 °C until they reached an OD_600_ of 0.6 to 0.7. Overexpression of the His-tagged proteins was induced with 0.5 mM IPTG at 15 °C. After incubation for 5 h at 15 °C to avoid the formation of inclusion bodies, the cells were collected by centrifugation, suspended in lysis buffer (20 mM Tris [pH 8.0], 500 mM NaCl, 1 mM β- mercaptoethanol), and disrupted using liquid nitrogen and a mortar. Cell debris were removed by centrifugation at 20,000 x g for 30 min at 4 °C, and the supernatant was loaded onto a nickel nitrilotriacetic acid (Ni-NTA)-agarose column (Invitrogen). The column was washed sequentially with lysis buffer containing 20 mM and 40 mM imidazole, respectively, and the His-tagged protein was eluted with the same buffer containing 250 mM imidazole (elution buffer). PD-Minitrap G-25 (GE Healthcare) columns were used to desalt and exchange the buffer to 20 mM Tris (pH 8.0) and 200 mM NaCl (working buffer). Vivaspin 500 (Sartorius) centrifugal filters (cutoff of 10 kDa) were used to concentrate the purified proteins. The purity of the overexpressed His-tagged protein was confirmed by sodium dodecyl sulfate (SDS)-PAGE 16% gels.

### Sequence analyses

Protein sequence alignments were performed using CLUSTALW (https://www.genome.jp/tools-bin/clustalw), and the alignments were visualized with Jalview 2 [21]. AnBLUF65 and AnBLUF46 3D modeled structures were performed using the Swiss-Model workspace/GMQE [22] in a search for template mode. In all cases, the best fit corresponded to the BlsA PDB structures (6W6Z for the dark-adapted form and 6W72 for the light-adapted form). Models were visualized and handled with PyMOL (by Schrödinger). Ring 2.0 webserver [23] was used to explore the specific ππ stacking interactions in all protein models as well as other non-covalent interactions.

### Spectroscopic measurements

Absorption spectra were registered using an Ocean Optics modular UV-Vis spectrophotometer USB2000+. The assays were done using a 5x5 mm quartz cuvette (Hellma, Germany), placed in a Quantum Northwest FLASH 300 cuvette holder connected to a Peltier-based temperature controller, containing 250 µl of air-saturated protein solution in 20 mM TRIS, 200 mM NaCl, pH 8.0 buffer. Scattering effects on the absorption spectra were corrected using a|e – UV-Vis-IR Spectral Software 1.2, FluorTools (www.fluortools.com).

Light-adapted state for AnBLUFs (l-AnBLUF) was obtained by blue light irradiation of the dark-adapted form (d-AnBLUF) with a 1 W Royal Blue LED (Luxeon Star LEDs) emitting at 443 ± 20 nm. Absorbance changes followed at 510 nm (Δ*A*) vs time (*t*) were recorded during dark-light cycles at different temperatures. The light-adapted-state formation time, τ_lBLUF_, and the back-recovery time through thermal recovery to the dark-adapted state, τ_rec_, were determined using the exponential equations 1 and 2, respectively, where *A* is the pre-exponential factor and *ΔA_0_* is the initial absorbance:

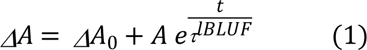

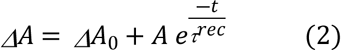

Fluorescence emission spectra were recorded with a Hitachi F-2500 spectrofluorometer equipped with a Hamamatsu R-928 photomultiplier. Neutral density filters (10 %T) were placed onto the excitation output beam to minimize photochemical processes during acquisition. Emission spectra were obtained by selective excitation of FAD cofactor at 460 nm. Temperature control was performed using a circulating fluid bath (Haake F3) connected to the cuvette holder.

### Protein extraction

*A. nosocomialis* was grown stagnantly in LB at 37 °C under blue light or in the dark for 24 hs, and the procedure was repeated to generate three independent biological replicates. The cells were recovered by centrifugation at 7,000 g for 10 min at 4 °C. The cells were then resuspended in 500 µl of extraction buffer (25 mM Tris pH 7.0), 0.5% Tween 20, 2 mM EDTA, 5 mM, β- mercaptoethanol) and lysed by sonication while keeping samples in ice. Cell debris and non lysed cells were collected by centrifugation at 12,000 rpm for 15’ at 4 °C and the supernatants were carefully decanted into clean tubes and stored at −80 °C. Total protein content present in the supernatant was quantified using bicinchoninic acid (BCA) (Thermo Scientific, Germany). Then, 30 µg total proteins were loaded in SDS-PAGE (10% stacking gel and 5% running gel), and allowed to separate electrophoretically only 1 cm long within the separation gel. The gel was incubated for at least 3 hs in fixing solution: 30% v/v ethanol, 2% H_3_PO_4_, and after being washed with MiliQ water, it was incubated for 1 h in staining solution: 18% v/v methanol, 17% p/v (NH_4_)_2_SO_4_ y 2% H_3_PO_4_ under vigorous shaking. Then, Coomassie G250 powder (0.5 g/L) was added and further incubated for 1 or 2 days until stained proteins were visible, which were then cut from the gel and sent to the Proteomics Core Facility of CEQUIBIEM at the University of Buenos Aires where protein digestion and Mass Spectrometry analysis were performed. Samples were resuspended in 50 mM (NH_4_)HCO_3_ pH 8.0, digested overnight with sequencing-grade modified trypsin (Promega) and desalted with Zip-Tip C18 (Merck Millipore). Proteins were analyzed by nanoHPLC (EASY-nLC 1000, Thermo Scientific, Germany) coupled to a mass spectrometer with Orbitrap technology (Q-Exactive with High Collision Dissociation cell and Orbitrap analyzer, Thermo Scientific, Germany). Peptide Ionization was performed by electrospray. Data were analyzed with Proteome Discoverer 2.1 software (Thermo Scientific, Germany) for identification and area quantitation of each protein. Protein identification was performed using *Acinetobacter nosocomialis* strain Ab1 protein collection as reference (UP000244598; https://www.uniprot.org/uniprot/?query=proteome:UP000244598) since a fully annotated *A. nosocomialis* RUH2624 proteome is not currently available. Missing value imputation method [24] was applied to the analyzed results, and Perseus software v1.6.1.3 (Max Planck Institute of Biochemistry) was used to perform the statistical tests. Proteins showing a fold change (FC) ≥ 1.5 between light and dark conditions and *p*-value < 0.05 were considered differentially produced.

### Proteomics Bioinformatics

Biological functional analyses of differentially over-represented proteins detected by proteomic approaches were categorized according to their molecular function, biological process, and cellular component, by using the Blast2GO tool.

## Results and Discussion

### Light modulates motility in a wide range of temperatures in *A. nosocomialis* RUH2624

We have recently shown that in *A. baumannii* ATCC 17978, light modulates motility from low to moderate temperatures up to 24 °C [11]. In this work, we assayed the effect of light on motility at different temperatures in *A. nosocomialis* RUH2624. Our results show that this microorganism grows at the inoculation point under blue light at 25 °C, and despite of moving increasingly as temperature raises, it only reaches 10-20 % of plate coverage at the highest temperatures such as 32 and 37 °C (Figs 1A and 1B). On the contrary, the bacteria moved covering the whole plates at all temperatures assayed in the dark (Figs 1A and 1B). Thus, in contrast to *A. baumannii*, in *A. nosocomialis* RUH2624 light modulates motility not only at environmental temperatures, but within a wide range of temperatures including those found in warm-blooded animals such as 37 °C.

**Fig 1.**
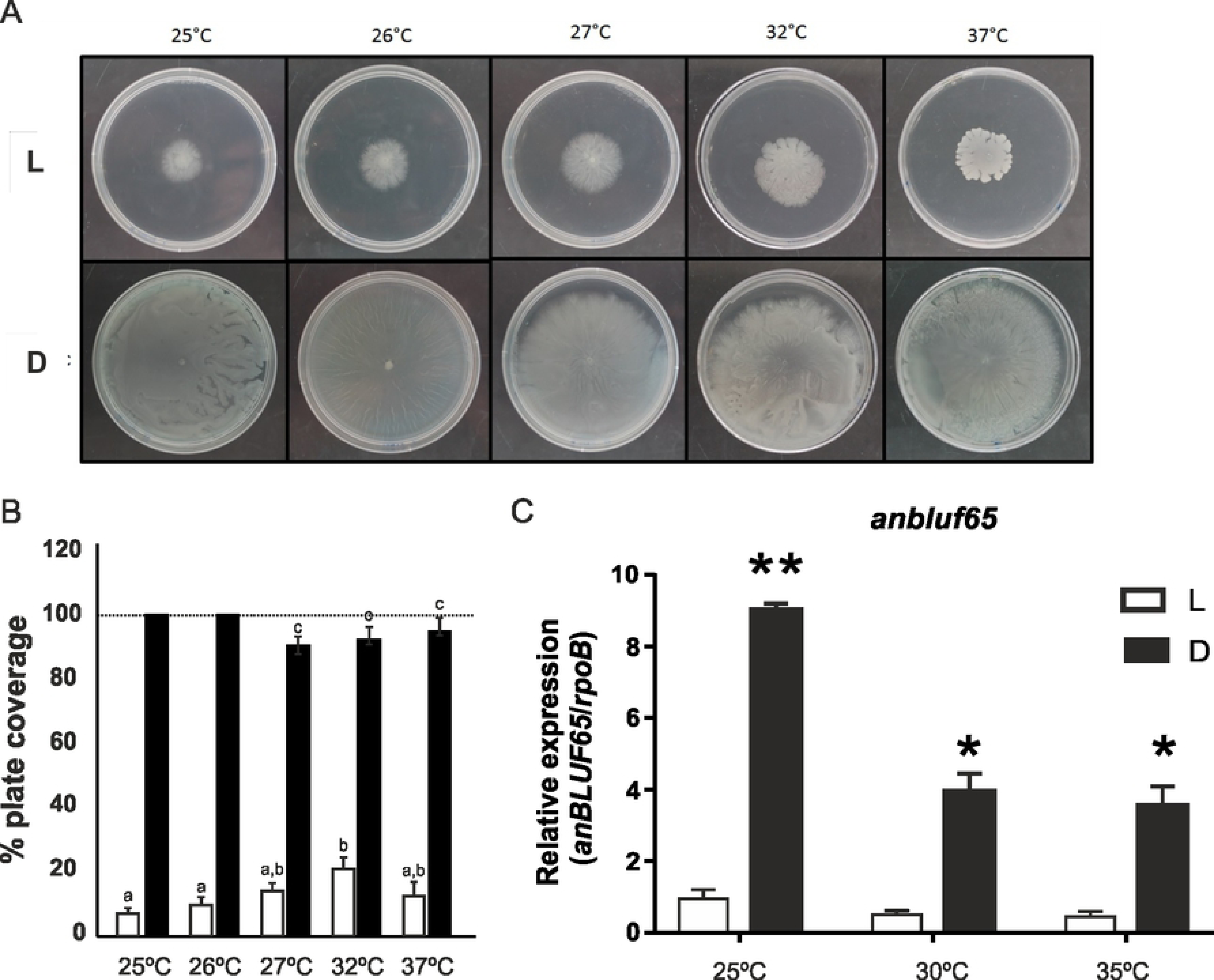
Effects of blue light and temperature on *A. nosocomialis* motility and *anbluf65* expression. (A) Cells of RUH2624 strain were inoculated on the surface of motility plates and incubated at the indicated temperatures. Plates were inspected and photographed after incubation in darkness (D) or in the presence of blue light (L) at the indicated temperatures. (B) Quantification of cell motility estimated as the percentage of plate coverage, i.e., the percentage of the Petri plate area covered with bacteria, in motility plates inoculated with RUH2624 wild type and incubated at the indicated temperatures. Three independent experiments were performed in each case. The area of plates covered with bacteria was measured with ImageJ (NIH), and then the percentage of plate coverage was calculated. The mean and standard deviation is informed. Different letters indicate significant differences as determined by ANOVA followed by Tukey’s multiple comparison test (p < 0.05). For those conditions at which the bacteria just reached the edge of the plate a value of 100% is informed. (C). cDNA from *A. nosocomialis* RUH2624 cells grown in motility plates in the presence of blue light (L) or in darkness (D) at different temperatures was used as template for quantitative real-time PCR using *anBLUF65* specific primers. Transcription of *rpoB* was used as a constitutively expressed internal control. The results are representative of three independent experiments. The mean and standard deviation are shown. *t*-test was used to determine whether two values were significantly different comparing data within each temperature assayed. *p*-values: *, p < 0.01; **, p < 0.001.

### *A. nosocomialis* RUH2624 encodes three BLUF-type putative photoreceptors

Comparative sequence analyses have shown the presence of three BLUF domain-containing proteins encoded in the *A. nosocomialis* RUH2624 genome [8]. These putative photoreceptors received accession numbers EEW98085, EEX00046, and EEW01065 in the Genbank database and will be further referred to here as AnBLUF46, AnBLUF65 and AnBLUF85, respectively .

Sequence alignments of these three BLUF-domains containing proteins with other well characterized BLUF domains such as Slr1694 from *Synechocystis* sp., Tll0078 from *Thermosynechococcus elongatus*, AppA and BlrB from *Rhodobacter sphaeroides,* and BlsA from *A. baumannii*, confirmed the presence of 22 highly conserved residues in the N-terminus of canonical BLUF domains according to pfam04940 (Fig 2A). Several BLUF domains belonging to different species from the genus *Acinetobacter* have been described so far as functional blue light photoreceptors [10–12, 15]. Alignment of these functional *Acinetobacter* BLUF-photoreceptor sequences show a higher level of conserved residues in their N-terminus domain (33 residues, Fig 2B), while the analysis of all γ-*Proteobacteria*, including 30 *Acinetobacter* sequences deposited in pfam04940 displays lower conservation (14 residues, Fig 2C) in their primary structure and in some cases, insertions between the conserved Tyr and Gln (Fig 2A), though with a low occupancy level (Fig 2B).

**Fig 2.**
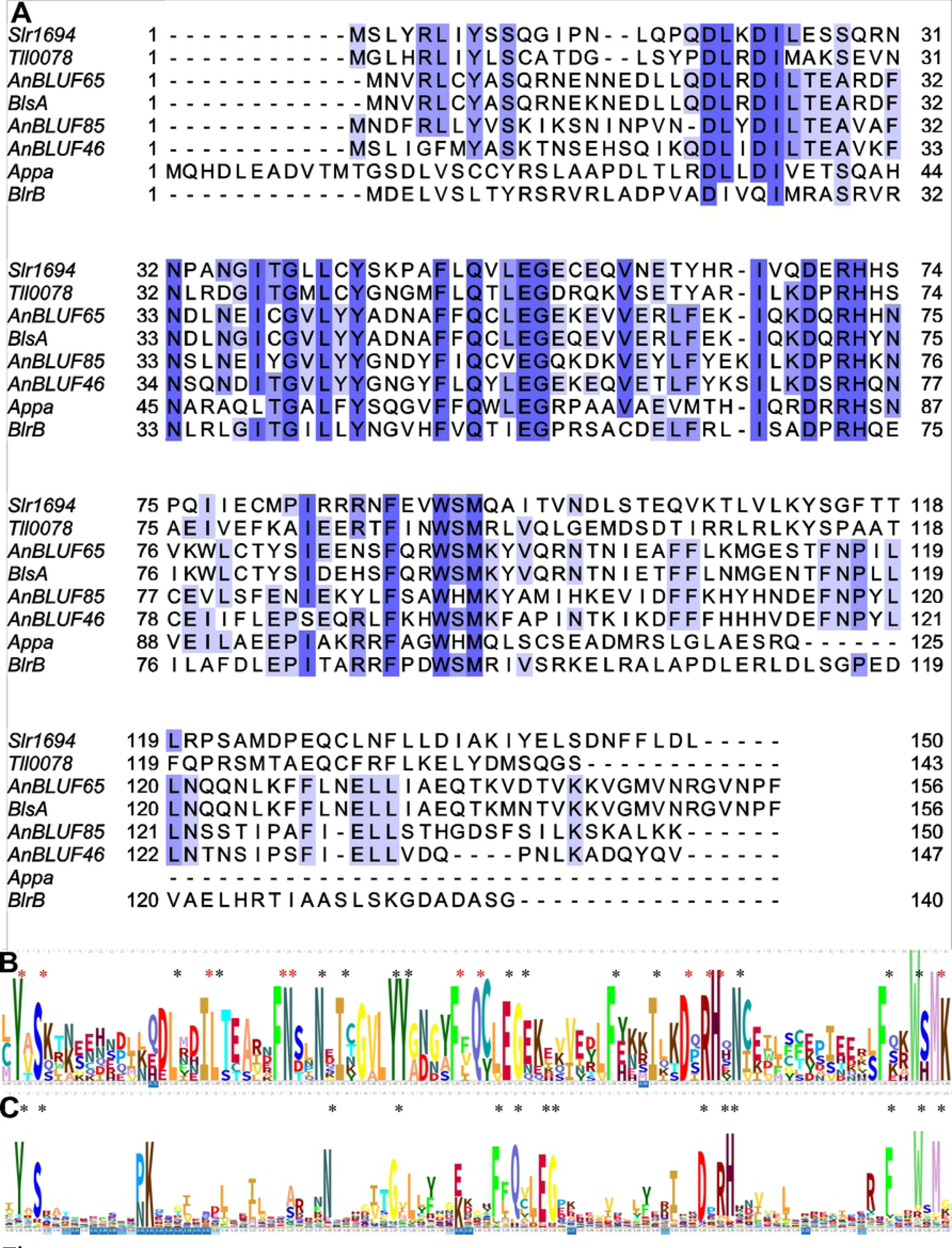
Comparative analysis of BLUF domains. (A). Multiple alignment of structurally characterized BLUF domains members of pfam04940 generated with ClustalW and adjusted with Jalview. Residues are colored according to the percentage of identity and the conservation level (darker, highly conserved). Asterisks show residues with high or very high conservation. (B) Alignment logo of *Acinetobacter* BLUFs directly or undirectly characterized as photoactives, i.e. BlsA (*A. baumannii*); AnBLUF65 (156 amino acids), AnBLUF 46 (147 amino acids) and AnBLUF85 (150 amino acids); Q7BC36, Q6FAI1 and Q6FAH9 (*A. baylyi* ADP1).(C) Alignment logo performed with 341 BLUF sequences from 204 γ- *Protebacteria* species, obtained from PF04940 (Pfam). Asterisks show residues with high or very high conservation interacting with flavin (red) or not (black).

Phylogenetic analyses indicate that AnBLUF65 and *A. baumannii*’s BlsA BLUF domains are grouped in a monophyletic cluster [8], thus indicating that they are orthologs. *anbluf65*’s genomic context shares many features with *blsA*’s (Fig 3). *anbluf65*’s 5’ upstream region is highly conserved with respect to the 5’ upstream region of BlsA in *A. baumannii* ATCC 17978 (Fig 3), as well as with different strains of *A. nosocomialis* (S1 Fig). Interestingly, the presence of insertion sequence ISAba27 was detected in the intergenic region upstream of the Bof coding sequence in *A. nosocomialis* RUH2624 (Fig 3A and S1 Fig). Whether ISAba27 is providing a hybrid promoter or disrupting the transcription of a putative operon that includes *anbluf65* is a possibility that deserves further experimentation. *A. nosocomialis* RUH2624 *anbluf65* 3’ downstream region is more variable compared both with BlsA or with other strains of *A. nosocomialis* (Fig 3 and S1 Fig). Gene *anbluf46* is located in a different genome location, surrounded by genes coding for a putative succinyl-CoA:3-ketoacid-coenzyme A transferase, a LysR transcriptional regulator, a hypothetical protein, a putative heat shock, a conjugal protein, and another hypothetical protein on the 5’ upstream region (Fig 3A). On the 3’ downstream region there are encoded: a putative protein with a metallo-β-lactamase fold, an H^+^/gluconate symporter, a hydroxybutyrate dehydrogenase, an AraC transcriptional regulator, and an aromatic acid/H^+^ symporter (Fig 3A).

**Fig 3.**
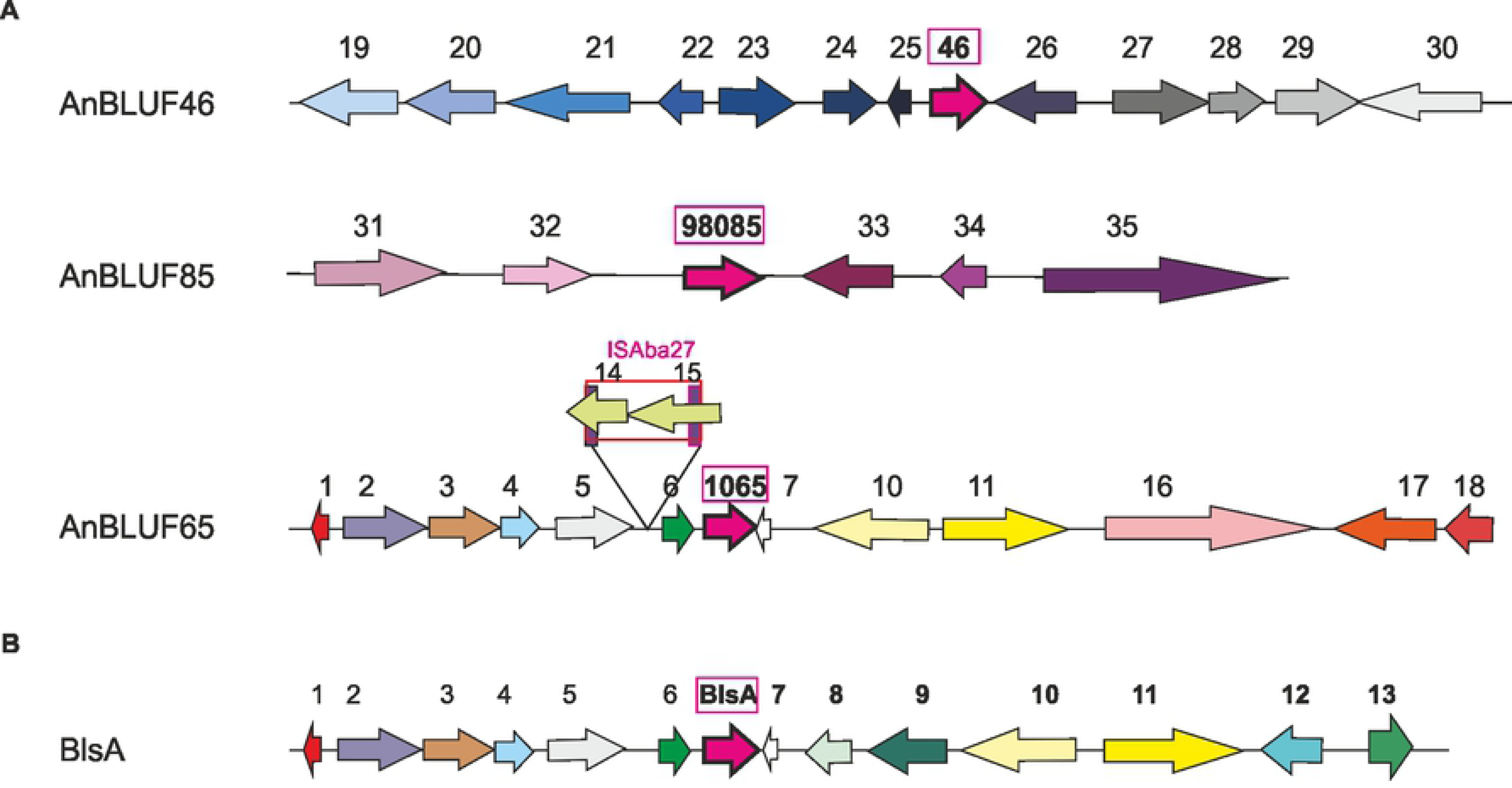
Genomic context of the three BLUF photoreceptors present in *A. nosocomialis* and comparison with BlsA of *A. baumannii*. (A). AnBLUF46, AnBLUF85 and AnBLUF65 genetic surroundings in *A. nosocomialis* RUH2624 strain. (B). BlsA genetic surroundings in *A. baumannii* strain ATCC 17978. Coding-sequences (CDSs) are located in their corresponding frame. The different photoreceptors are indicated as pink arrows. Different colors in CDSs indicate different functions. Gene annotations are indicated as numbers above the schemes, with the following codes: 1-DUF2171; 2-Acyl-CoA dehydrogenase; 3-GlcNAc-PI de-N-acetylase; 4- Methyltransferase domain; 5-Glycosyltransferase; 6-BOF- Class 2b aminoacyl-tRNA synthetases--NirD/ YgiW/YdeI family stress tolerance protein; 7-HP; 8-RhtB - Homoserine/Threonine/Homoserine Lactona translocator; 9-AraC transcriptional regulator; 10- Poly(R)-hydroxyalkanoic acid synthase; 11-GltS- Sodium/glutamate simporter; 12- DUF815; 13-FdaR- Fatty acid transcriptional regulator; 14-DDE endonuclease domain, putative transposase; 15-Helix-turn-helix of DDE superfamily endonuclease; 16- Proton antiporter-2 (CPA2) family; 17-NDAB-Rossmann Superfamily- Oxidoreductase; 18-DoxX-like family; 19-Succinyl-CoA:3-ketoacid-coenzyme A transferase subunit B; 20-Succinyl-CoA:3-ketoacid-coenzyme A transferase subunit A; 21-LysR family transcriptional regulator; 22- HP; 23-DUF3298-Putative heat-shock protein; 24-DUF2846-Putative conjugal domain; 25-HP; 26-MBL fold metallo-hydrolase; 27-H+/gluconate symporter; 28-3-Hydroxybutyrate dehydrogenase; 29-AraC family transcriptional regulator-RmlC-cupin protein; 30-Aromatic acid:H+ symporter (AAHS); 31-Conjugal transfer pilus assembly protein TraB; 32-N- Acetyltransferase; 33-HP; 34-FAM199X; 35-Relaxase-MobA/MobL family protein. HP: hypothetical protein.

*anbluf46* does not seem to be part of an operon, as both flanking genes are encoded in the opposite direction (Fig 3A). Comparing *anbluf46* genomic context in strain RUH2624 with other strains of the same species, it is observed that the 3’ downstream region is highly conserved while the 5’ upstream region is variable, showing the presence of transposases in different locations (S2 Fig). *anbluf85* was only found encoded in *A. nosocomialis* RUH2624 and UBA873 strains (100% identity). In RUH2624, *anbluf85* is flanked by a 5’ upstream region coding for an acetyltransferase and a putative conjugal transfer pilus assembly protein TraB. On the 3’ downstream region there are encoded a hypothetical protein, a FAM199X domain-containing protein, and a relaxase MobA/MobL (Fig 3A). RUH2624 *anbluf85* genomic context is different from that in UBA873 (S3 Fig). In the latter, only the 3’ downstream region is shown as the contigs are not assembled yet (S3 Fig). Sequence comparisons indicate the presence of an *anbluf85* homolog in *Acinetobacter lwoffii* strain M2a plasmid pAVAci116 as well as in *Acinetobacter* sp. ACNIH1 plasmid pACI-148e with 100% identity at nucleotide level. Interestingly, also an *anbluf85* homolog with 98.13% identity at the nucleotide level was reported to be present in *Acinetobacter baumannii* isolate KAR [25]. Moreover, 100% identity homologs at the aminoacidic level are also present in *A. baumannii* strains A30, A10, and MSHR_A189, two in *A. radioresistens* strains 50v1 and TG29429; and in *A. baumannii* strains MRSN7353 and ARLG1306 showing 1 aminoacidic difference with AnBLUF85. Thus *anbluf85* is not widely distributed but its presence is observed discretionally in some *Acinetobacter* species, and the possibility of horizontal gene transfer is suggested.

### Only AnBLUF65 and AnBLUF46 are stable photoreceptors

We evaluated next whether the three BLUF-domain containing genes encode active photoreceptors. For this purpose, genes *anbluf65*, *anbluf46* and *anbluf85* were recombinantly expressed and purified to assess their functionality.

Fig 4 shows the UV-vis spectra of AnBLUF65 (92% identity with BlsA) and AnBLUF46 (45% identity with BlsA) at 15 °C in working buffer. Before blue light irradiation, both dark-adapted states of AnBLUF65 (d-AnBLUF65) and AnBLUF46 (d-AnBLUF46) showed the typical absorption band of the fully oxidized flavin corresponding to the transition S_0_→S_1_ with λ_max_ ≈ 460 nm and shoulder approximately at 483 nm. Upon blue-LED illumination, the band is red-shifted, characteristic of the formation of the light-adapted state of BLUF proteins (l-BLUF) [26], with absorption maximum wavelength shift of 8 and 12 nm for l-AnBLUF65 and l-AnBLUF46, respectively (Fig 4A and C). The differential absorption spectrum (Δ*A*) between the light- and dark-adapted states of AnBLUF65 showed positive changes at 508 and 472 nm, and negative changes at 455 and 427 nm. AnBLUF46 also showed positive changes at 511 and 476 nm, whereas negative bands were less defined (insets Fig 4A and C). The overall results indicate that AnBLUF65 and AnBLUF46 are active photoreceptors. Gene *anbluf85* was weakly expressed and the partially purified product was barely stable precipitating at T higher than 12 °C, with a photocycle presenting a ≈ 7 nm red shift (S4 Fig).

**Fig 4.**
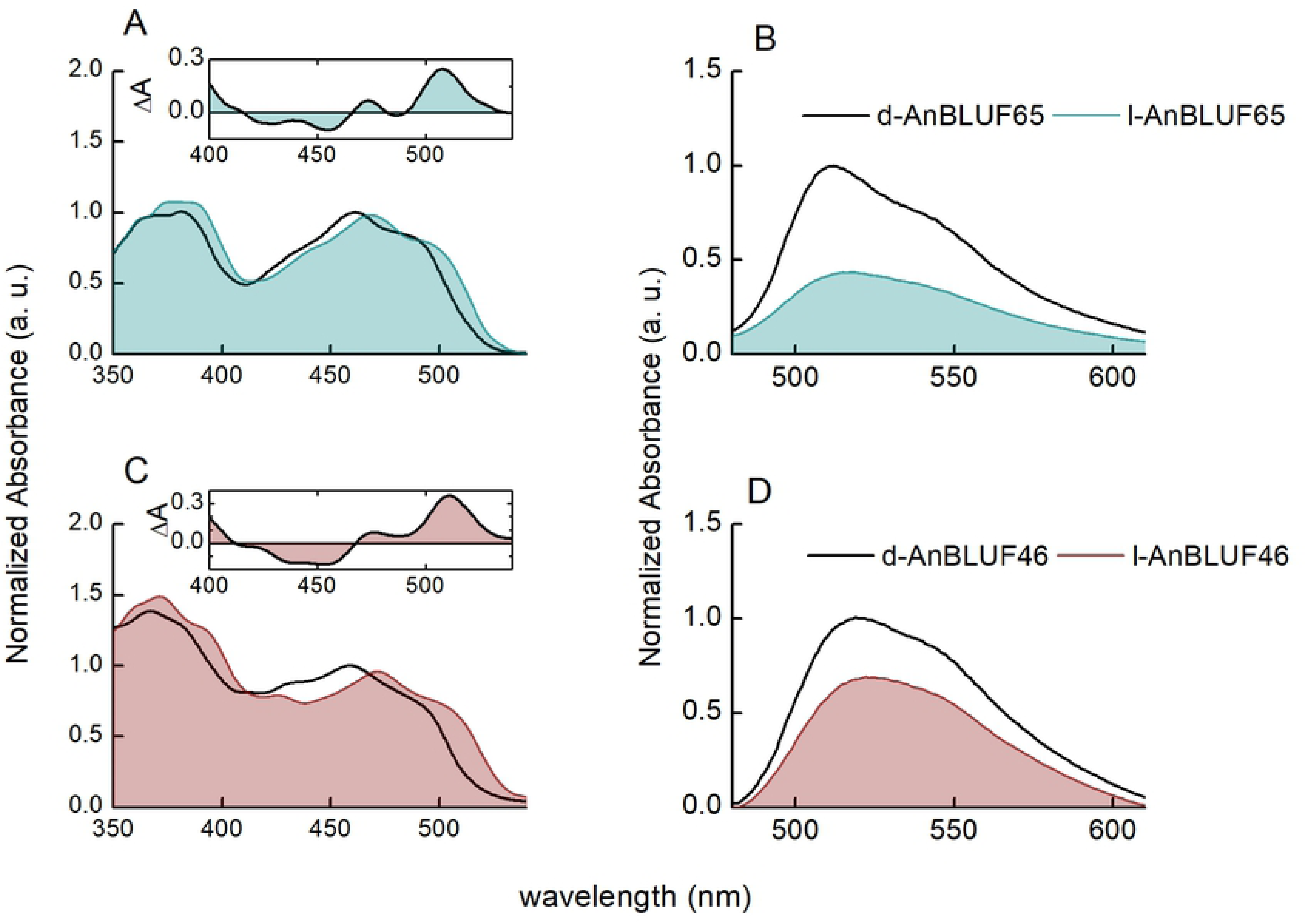
Normalized steady-state visible absorption-emission spectra changes of photoreceptors by blue light illumination, in working buffer solution at 15 °C. (A and B). dAnBLUF65 (black line) and lAnBLUF 65 (blue line). (C and D). dAnBLUF46 (black line) and lAnBLUF46 (red line). Inset: differential absorption spectra (lAnBLUF-dAnBLUF).

Fluorescence emission spectra obtained by excitation at 460 nm at 15 °C for d-AnBLUF65 and d- AnBLUF46 also depicted a BLUF-like behavior, with an emission maximum at 512 and 519 nm, respectively (Fig 4B and D). The fluorescence bands of both proteins were blue-shifted compared with free FAD in the same working buffer [10, 11] confirming the existence of specific interactions between the isoalloxazine group of the flavin with the surrounding residues, and reducing solvent accessibility within the flavin binding pocket [27]. When the fluorescence spectrum of the light-adapted and the dark-adapted forms of both photoreceptors were compared, they exhibited the typical red shift of approximately 5 nm of the fluorescence maximum in addition to a loss of vibratory structure and a lower emission intensity, suggesting a more relaxed environment around the cofactor [28–30] [10].

### Only *anbluf1065* is expressed in *A. nosocomialis* RUH2624

The expression levels of the three BLUF-type photoreceptors encoded in *A. nosocomialis* RUH2624 were analyzed on cells recovered from motility plates incubated at a wide range of temperatures (25–35 °C) under blue light or in the dark by qRT-PCR as described previously [14]. We were able to detect expression of *anbluf1065* only, the *blsA* ortholog in *A. nosocomialis* [8]. *anbluf65* was expressed at 25, 30, and 35 °C, with approximately 9, 8, and 7.5-fold higher expression in the dark than under blue light (Fig 1C); a response previously observed for BlsA [11, 15]. Moreover, expression levels were higher at lower temperatures (Fig 1C). Thus, our results suggest that expression of the putative *anbluf65* photoreceptor correlates with regulation of motility by light.

### AnBLUF65’s photoreceptor activity is operative in a temperature range that correlates with regulation of motility by light

*A. nosocomialis* modulates motility in response to blue light in a similar fashion as observed for *A. baumannii*, but in a wider range of temperatures. Table 2 summarizes the temperature dependency of visible absorption and emission fluorescent properties of flavin in purified AnBLU65 and AnBLUF46 photoreceptors. Absorption maxima of the dark-adapted forms of both proteins were blue-shifted when temperature increased, reaching in the case of d-AnBLUF46 the value of free FAD in solution (≈ 450 nm). The ratio between the shoulder absorbance at ≈ 480 nm and the absorption maximum (A_sh_/A_max_) diminished with temperature increments, as did the redshift upon blue light illumination, suggesting a distancing between the conserved Tyr to the isoalloxazine ring [31]. Fluorescence emission properties of the cofactor were significantly affected upon formation of l-AnBLUF65 at 24 and 37 °C, with a redshift in response to blue light, as described before at 15 °C or when temperature increased, implying a relaxation in the hydrogen bond network surrounding the isoalloxazine, and/or changes in the polarity in the chromophore cavity as a consequence of increased solvent accessibility. Widening of the emission spectrum full-width half-maximum (FWHM) in response to the formation of the light-adapted state for AnBLUF65 was observed at 24 and 37 °C, and this effect was also evident when temperature increased. Overall, these results indicate that the flavin environment in AnBLUF65 is affected by temperature, as it has been shown for BlsA [10, 11]. In contrast, emission maximum wavelength and FWHM for the dark- or the light-adapted states of AnBLUF46 did not show significant changes neither in response to blue light at 24 and 32 °C, nor to temperature increments. In fact, d- and l- AnBLUF46 isoalloxazine pockets appeared to be more polar than the same states of AnBLUF65. Chromophore environments in AnBLUF65 and AnBLUF46 were somewhat dissimilar and appeared to react in a different way to the formation of the light-adapted state. This fact goes in good agreement with structural predictions from models of these two proteins generated using the crystallographic structure of dark- and light-adapted states of BlsA (PDBs 6W6Z and 6W72, respectively) as templates (Fig 5). Fig 5A and 5B show the location of water molecules (blue dots) for the dark-adapted form of BlsA (d-BlsA) and the light-adapted (l-BlsA) respectively, determined by Chitrakar *et al.* [32]. The flavin pocket access site has more affinity and/or accessibility to water molecules in the light-adapted form, supporting our previous characterization of chromophore behavior inside BlsA cavity by the flavin fluorescence emission (Fig 5A and B)[10]. Modeled structures of AnBLUF65 (Fig 5C and 5D), which is BlsA ortholog as described before, show that the environment of the cofactor is conserved compared with BlsA. This means that analogously to BlsA, the flavin pocket in the light-adapted form of AnBLUF65 is more solvent-accessible. In contrast, the solvent accessibility to the chromophore cavity between d- and l-AnBLUF46 is indistinguishable (Fig 5E and F). In addition, the flavin binding site in AnBLUF46 seems more solvent-accessible than in BlsA/AnBLUF65. Hence, these data stress out how temperature and blue light differentially affect them.

**Fig 5.**
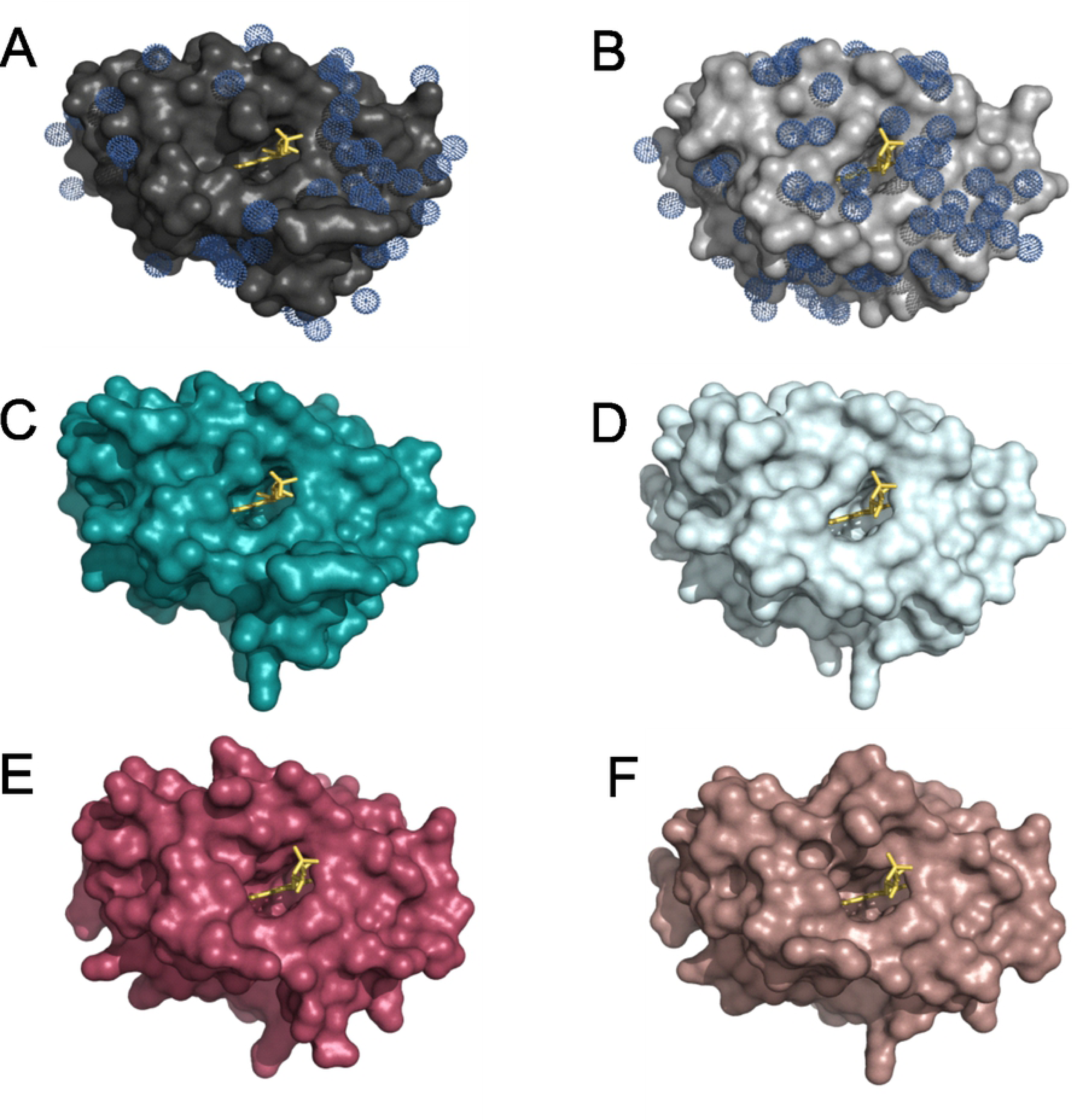
Surface representation of *Acinetobacter* BLUFs and FMN (yellow sticks) into the site access pocket. Internal surface associated with the cavity is also depicted. (A) BlsA dark-adapted, PDB 6W6Z; (B) BlsA light-adapted, PDB 6W72, (C) d-AnBLUF65; (D) l-AnBLUF65; (E) d-AnBLUF46 and (F) l-AnBLUF46.- Models were generated with the Swiss-Model server, using PDB 6W6Z as template for figure C and E and PDB 6W72 for figure D and F. Water molecules are depicted as blue dots.

**Table 2.** Absorption-emission properties of AnBLUF46 and AnBLUF65 at different temperatures.

|  | T<br>( °C) | Absorption |  |  | Emission |  |  |  |
| --- | --- | --- | --- | --- | --- | --- | --- | --- |
| | | $\lambda_{\max}$ | $A_{sh}/A_{\max}$ | $\Delta A$ | $\lambda_{\max}$ | d-AnBLUF | $\lambda_{\max}$ | l-AnBLUF |
|  |  | d-AnBLUF<br>(nm) | d-AnBLUF | (a.u) | d-AnBLUF<br>(nm) | FWHM<br>(nm) | lBLUF<br>(nm) | FWHM<br>(nm) |
| AnBLUF65 | 15 | 460 | 0.86 | 8 | 512 | 64 | 517 | 74 |
|  | 24 | 459 | 0.87 | 6 | 514 | 65 | 519 | 74 |
|  | 37 | 457 | 0.79 | 3 | 520 | 74 | 523 | 76 |
| **AnBLUF46** | 15 | 459 | 0.81 | 13 | 519 | 68 | 522 | 68 |
|  | 24 | 457 | 0.81 | 13 | 523 | 75 | 523 | 75 |
|  | 32 | 450 | 0.72 | 8 | 523 | 75 | 523 | 76 |

Maximum absorption wavelength ( $\lambda_{\max}$ ), shoulder to maximum absorbance ratio ( $A_{\text{sh}}/A_{\max}$ ), redshift maximum upon BL illumination ( $\Delta\lambda$ ), maximum emission wavelength ( $\lambda_{\text{em}}$ ) and full-width at half maximum (FWHM) for dark and light-adapted proteins.

Insets in Fig 4 show difference spectra both for AnBLUF65 and AnBLUF46 (lAnBLUF-dAnBLUF). In previous work, we have used the ΔA at 510 nm at different temperatures to follow the kinetics of formation of the light-adapted state of each protein, and in darkness, the recovery time back to the dark-adapted state, through cycles of blue light illumination. This kinetic profile was used to determine the quantum yield of the protein light-adapted formation, Φ_lAnBLUF_. Fig 6 presents the kinetic profiles of AnBLUF65 (Figures 6A and 6B) and AnBLUF46 (Fig 6C and 6D) tested at two temperatures. For AnBLUF65 the kinetic curves were monitored at 24 °C and 37 °C, which correspond to environmental and warm-blooded host temperatures, respectively. However, AnBLUF46 was tested at 24 °C and 32 °C, the maximum temperature at which this photoreceptor presented activity. Table 3 summarizes the light-adapted-state formation time τ_lAnBLUF_, and the recovery time back to the dark-adapted state τ_rec_, for both BLUFs as a function of temperature along with the photoactivation quantum yield of light-adapted state, Φ_lAnBLUF_.

**Fig 6.**
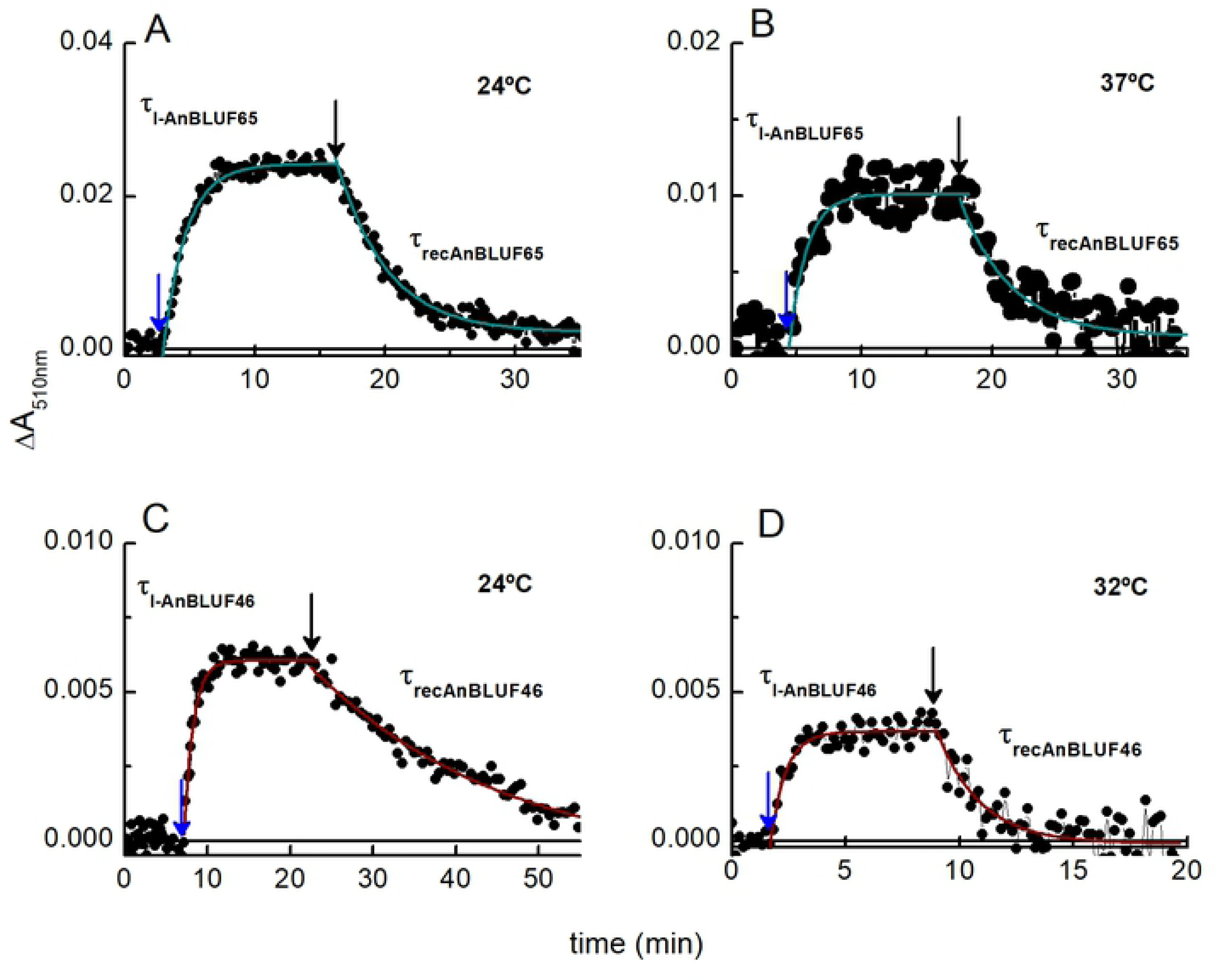
Kinetic profile of absorbance changes at 510 nm. (A and B) AnBLUF65 at 24 and 37 °C. (C and D) AnBLUF46 at 24 and 32 °C. Dark adapted protein, dAnBLUF was illuminated using blue light LED at 443 ± 20 nm (blue arrows). After maximal conversion to lAnBLUF, blue light was turned off (black arrow) and protein back to dAnBLUF.

**Table 3.** Kinetic parameters observed for AnBLUF65, AnBLUF46, and BlsA at 24 °C and 37 °C.

| | Temperature<br>( °C) | $\tau_{I\text{-BLUF}}$<br>(s) | $\Phi_{I\text{-BLUF}}$ | $\tau_{\text{rec}}$<br>(s) |
| --- | --- | --- | --- | --- |
| AnBLUF65 | 24 | 106±8 | 0.36±0.04 | 233±8 |
|  | 37 | 80±15 | 0.12±0.03 | 240±60 |
| AnBLUF46 | 24 | 70±4 | 0.11±0.01 | 950±90 |
|  | 32 | 40±5 | 0.08±0.02 | 115±20 |
|  | 37 | ND | ND | ND |
| BlsA* | 24 | 70±2 | 0.17±0.01 | 111±10 |
|  | 26 | 28±9 | 0.11±0.02 | 40±9 |
|  | 37 | ND | ND | ND |

ND: not detectable. \* [11]

Temperature increments had a significant effect on the *τ*_lAnBLUF_ for both proteins, with reductions of 25 and 40% for *τ*_lAnBLUF65_ and *τ*_lAnBLUF46_, respectively. AnBLUF46 had similar values to BlsA at 24 °C (Table 3), however, *τ*_lAnBLUF65_ was larger than that of BlsA. Analyses of *τ*_rec_ show that AnBLUF46 takes 15 mins to recover, nine times slower than BlsA; while AnBLUF65 was only two-fold slower than the *A. baumannii* protein. As expected for a thermal process, *τ*_recAnBLUF46_ was nine times faster when temperature increased to 32 °C, becoming similar to the value of *τ*_recBlsA_ at 24 °C. However, this was not the case of AnBLUF65, whose *τ*_recAnBLUF65_ remained constant at 24 and 37 °C. This behavior prompts us to speculate whether this recovery time might have some physiological meaning since AnBLUF65 presents conformational changes at least in its binding cavity upon illumination, but the time to return to the dark form is conserved.

Despite the high sequence similarity between AnBLUF65 and BlsA (92%), the behavior of the former was significantly different compared with that of the latter, regarding the efficiency to respond to blue light generating the light-adapted form. In fact, ϕ_lAnBLUF65_ was more than two-fold higher than that of BlsA at 24 °C, which can be interpreted as AnBLUF65 being more efficient to form the light-adapted state. The *τ*_lAnBLUF65_ was also significantly higher than that for BlsA, although to a lesser extent, and the *τ*_recAnBLUF65_ was two-fold higher than *τ*_recBlsA_, probably because the adapted form is more stabilized in AnBLUF65 than in BlsA. Another contrasting effect between such similar proteins is that AnBLUF65 is less prone to aggregate upon temperature increments than BlsA, which shows this effect macroscopically above 30 °C, while AnBLUF65 does not aggregate at 37 °C showing only incipient turbidity in the buffer solution. While ϕ_lBlsA_ reaches non- detectable values at T higher than 28 °C, ϕ_lAnBLUF65_ at 37 °C is 0.12±0.03. In contrast, ϕ_lAnBLUF46_ was not significantly impacted between 24 °C and 32 °C. Nevertheless, AnBLUF46 lost all photoactivity at T>32 °C. These results suggest that AnBLUF65 is more efficient to form the light-adapted state than AnBLUF46 at 24 °C.

### Intramolecular interactions and their role in intra-protein signaling

So far, we have analyzed these photoreceptors describing intrinsic phenomena occurring mostly in the N-terminal side of the protein, where the flavin resides in its cavity. These data have profiled AnBLUF65 and AnBLUF46 in the first part of the photo-reception process. Next, we aimed to expand our analyses to cover the subtle structural rearrangements between the dark and the light-adapted states of the protein. Intra-protein changes in hydrophobic and aromatic residue interactions driven upon illumination have been suggested by Chitrakar et al, 2020 to explain how the photo-signal might be translated from the flavin surroundings to the variable domain. Based on the similarity between BlsA and AnBLUF65, we modeled structures for both *A. nosocomialis* proteins, AnBLUF65 and AnBLUF46 (Fig 5), using BlsA as template, which is appropriate since BLUF’s in *Acinetobacter* genus share high sequence similarity. We used RING 2.0 [23] to profile all intramolecular interactions, such us π-π stacking, H-bonding, Van der Waals (VDW) interactions, and salt bridges, in AnBLUF65 and AnBLUF46, as presented in S1 Table. The analysis of π-π stacking interactions amongst aromatic residues helped us to profile aromatic networks for both proteins, in both dark- and light-adapted forms.

The first aromatic cluster found surrounds the flavin cofactor and involves the hydrophobic side of the isoalloxazine ring (ring I), the conserved Tyr, a Phe and a His for AnBLUF65 as shown in Fig 7. This His belongs to the already described motif Asp-X-Arg-His and it has been suggested that the protonation/deprotonation of the conserved Gln via His is energetically favourable [33]. In AnBLUF65, His73 forms π-π stacks with Tyr7 and with Phe49 in addition to other interactions i.e. H-bonds that have been described before. After blue light illumination this cluster does not show a significant impact on its Cα main chain or in the angle of the aromatics involved. This might be necessary to conserve the hydrophobicity in the flavin pocket [34].

**Fig 7.**
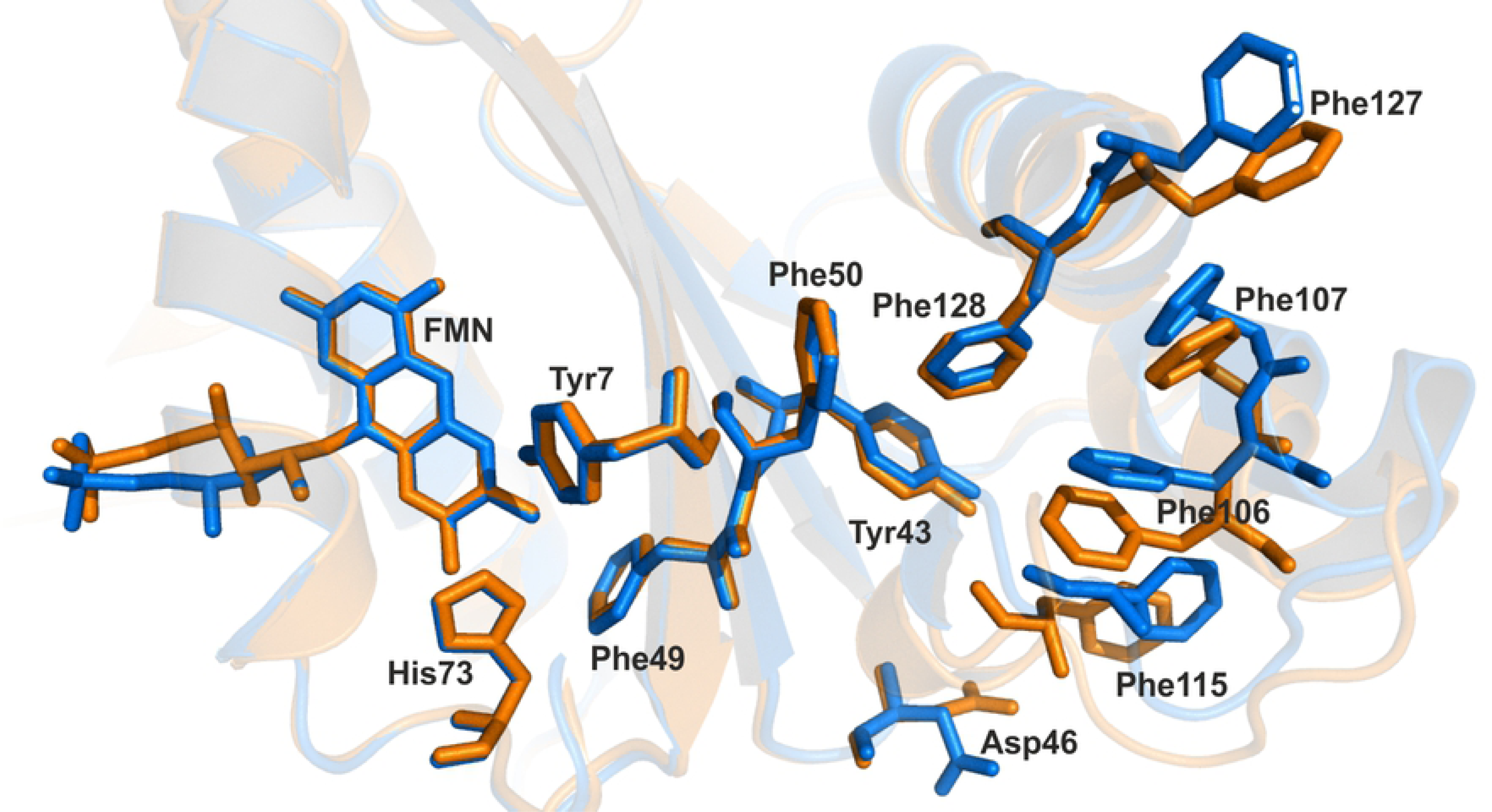
Residues involved in π-π stacking interactions in AnBLUF65 dark-(orange) and light-adapted (blue) state.

The next aromatic network found is in the variable domain (C-terminal) and contains two aromatic tetrads, that are formed by six Phe in AnBLUF65 (Fig 7). Upon blue light illumination, distances between aromatic centroids in these tetrads are shortened in all cases in AnBLUF65. Phe106 loses its π-π stack interaction with Phe115, being left only with VDW interactions. Thus, the formation of the light adapted state of AnBLUF65 requires a local re-arrangement of α-helices in the C-terminal and the aromatic network in this area is strengthened.

A Tyr-Tyr motif, highly conserved in *Acinetobacter* BLUFs (Fig 2B) also participates in the aromatic networks described before. AnBLUF65 has Tyr44 interacting with Phe49 on the chromophore side of the protein, while Tyr43 interacts through π-π stacking with Phe128 on the other side of the β- sheet, towards the variable domain (Fig 7). The notable re-arrangement is found towards the variable domain may be possible beacause an Asp residue located in the loop between the Tyr-Tyr motif and the first Phe-Phe motif, rotates upon blue light illumination, losing its contact with Phe115. Thus, one way to explain these findings is that the aromatic clusters found in the variable domain might internally stabilize the structure of this protein upon the formation of the light-adapted state. In contrast, the aromatic cluster in the flavin/N-terminal side of these proteins does not suffer any alteration when changing states. Taken together, it suggests that intramolecular hydrophobic/aromatic interactions might have a role in the elasticity of the protein monomer to go back and forth between the two states.

### Quantitative differential proteomic profiling in response to light in ***A. nosocomialis* RUH2624 at 37 °C.**

In previous studies, we demonstrated that light can modulate important physiological traits related to bacterial physiology and virulence in *A. baumannii*, *S. aureus*, *P. aeruginosa* and *A. nosocomialis* at temperatures found in warm-blooded hosts [8, 9, 14, 15, 17, 18, 35]. To broaden our knowledge of the response to light in *A. nosocomialis* at 37 °C, a quantitative comparative proteomic analysis was conducted to obtain an overall representation of the total cellular changes that occur in RUH2624 cells at the protein level upon blue light illumination at the mentioned temperature. We aimed to use proteomics as a tool to identify those proteins that could be present in different amounts under light and dark conditions and how these proteins could affect *A. nosocomialis’* physiology.

In total, 38 proteins were over-represented (fold change (FC) > 1.5 and *p-value* < 0.05, see Tables 4 and 5) when cells were grown under blue light or dark state at 37 °C (Fig 8A). Twenty proteins were accumulated in higher amounts in cells cultured under blue light with respect to darkness (Fig 8A and B), while eighteen proteins were found to be in a greater amount under dark state (Fig 8A and C). When categorized using BLAST2GO suite, these proteins were dispersed across a wide variety of functions, according to gene ontology terms (Fig 9). In the “Biological Process” and “Molecular Function” sections the majority of the over-represented proteins fell into the metabolic/cellular process and catalytic activity categories, respectively, indicating changes in the metabolism of this bacterium in response to light. Considering “Cellular Component”, over- represented proteins under light conditions appear mainly distributed in the plasma and outer membranes, while in darkness most of them are cytoplasmatic (Tables 4 and 5).

**Fig 8.**
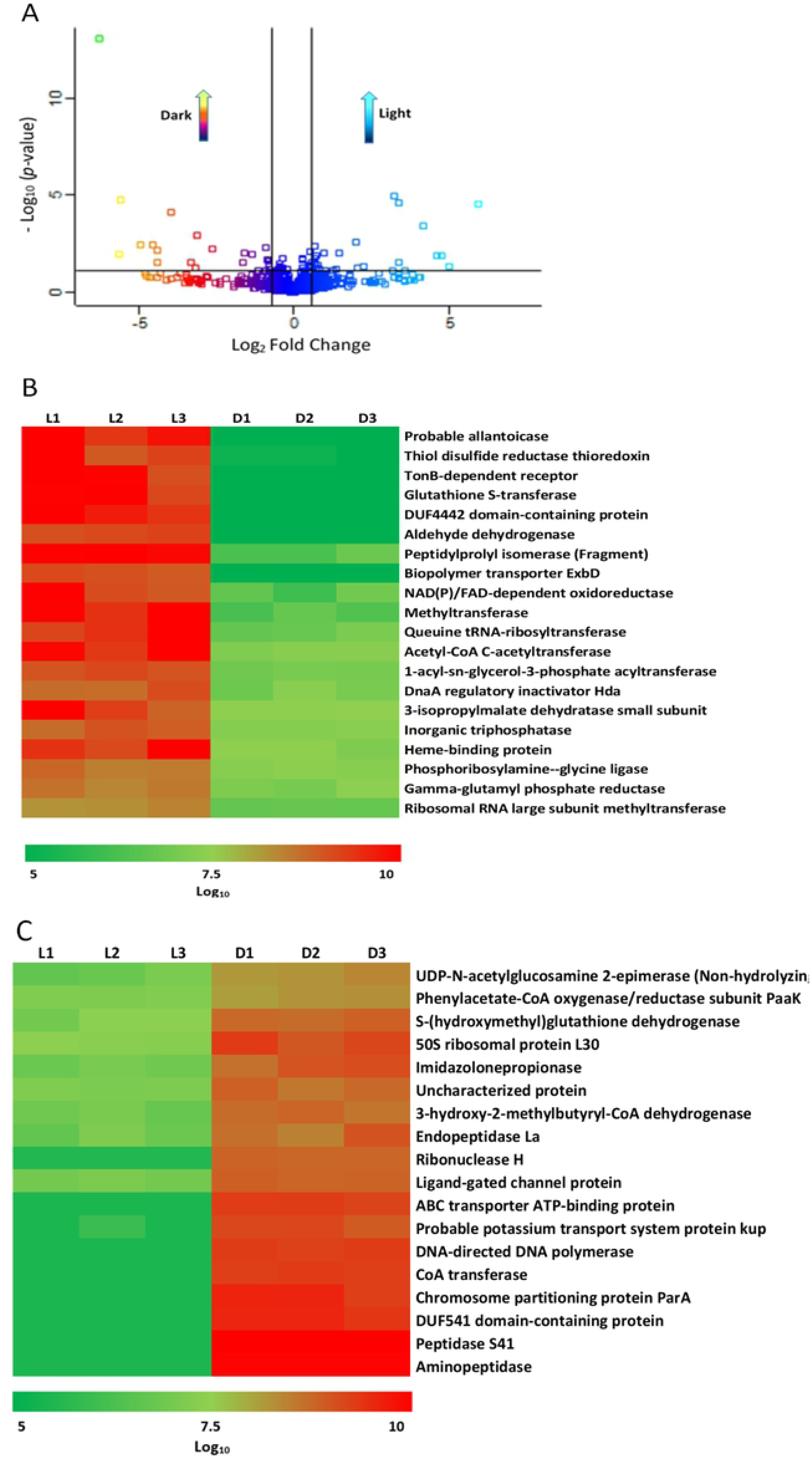
Comparison of over-represented proteins detected by nanoHPLC under light and dark conditions for *A. nosocomialis* RUH2624. (A) Scatter plot. (B) Heatmap of proteins produced in higher amounts under blue light. (C) Heatmap of proteins produced in higher amounts under dark condition.

**Fig 9.**
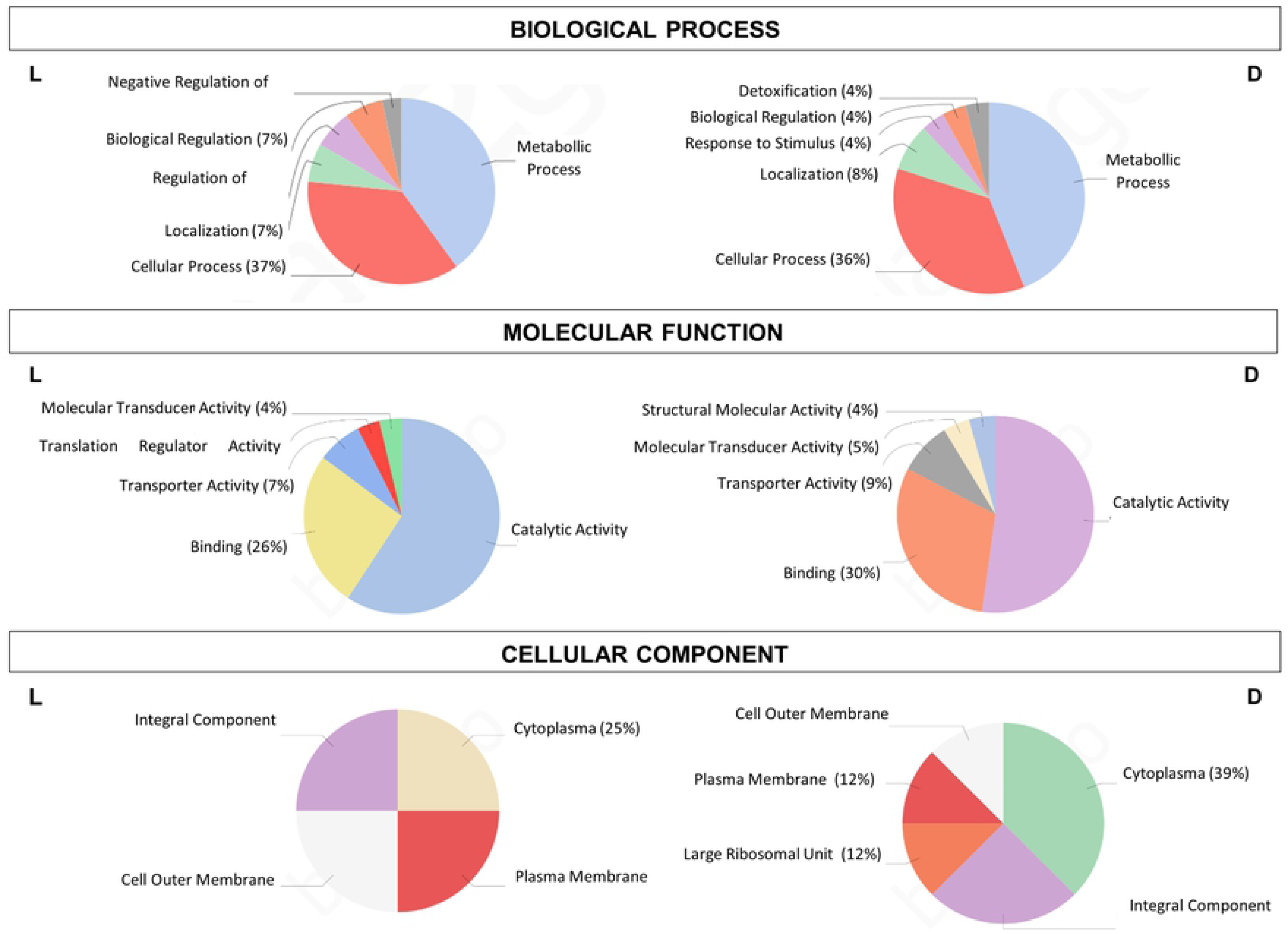
Gene ontology categories after Blast2Go analysis of all differential produced proteins. **L** and **D** correspond to over-represented proteins under light and dark conditions, respectively, for *A. nosocomialis* RUH2624.

**Table 4.**
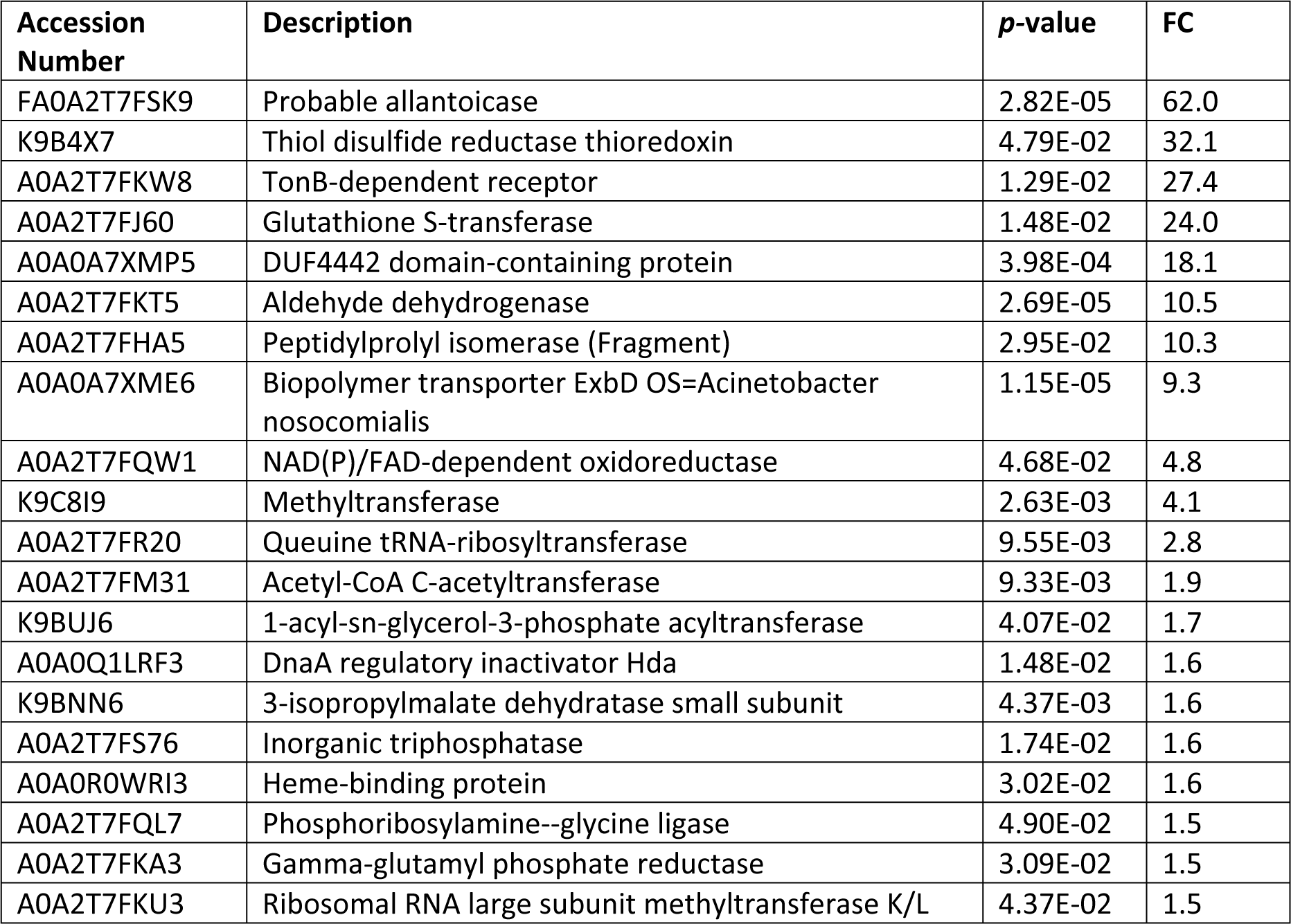
Proteins with increased abundance in the light compared to dark at 37 °C.

**Table 5.**
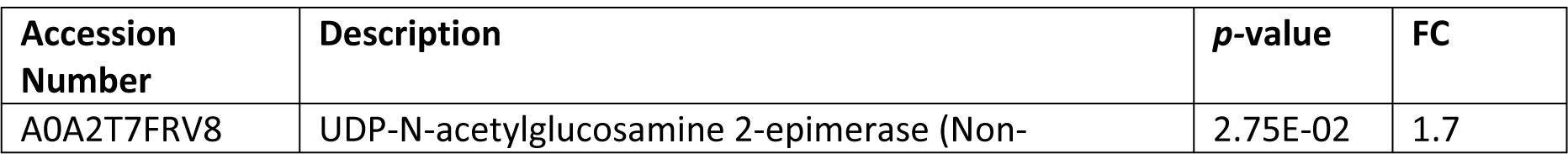
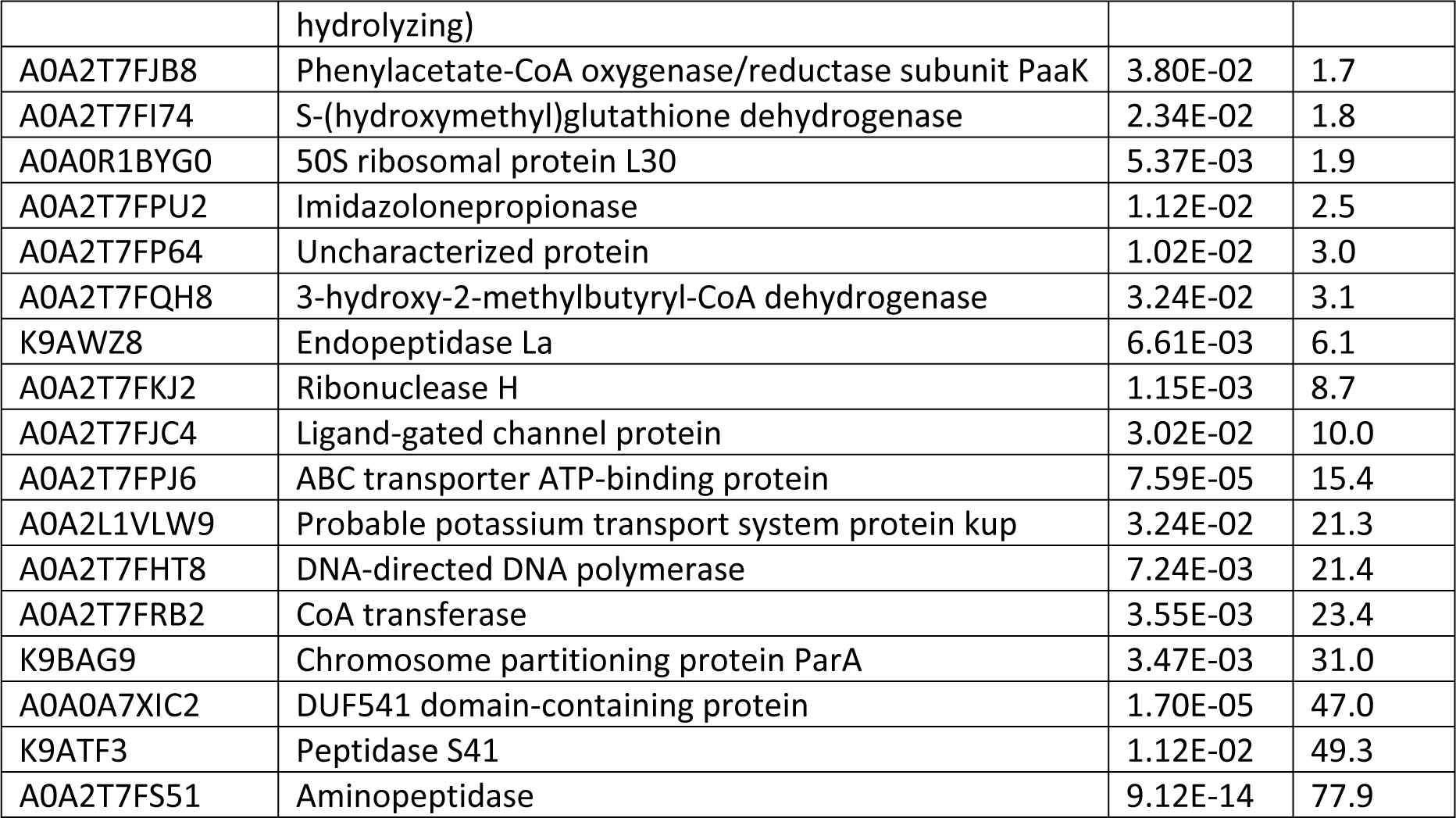
Proteins with increased abundance in dark compared to light at 37 °C

### Proteins with increased abundance in light compared to darkness

Light regulates the synthesis of proteins involved in a wide range of cellular functions. As mentioned before, these differentially over-represented proteins include someplaying roles in metabolism, stress responses, and virulence in different bacterial pathogens.

Two proteins, PurD and Alc, related to purine metabolism were over-represented (FC_PurD_= 1.50, FC_Alc_= 63.0) in the presence of blue light. PurD is a phosphoribosylamine-glycine ligase and participates in the synthesis of inosine 5’-monophosphate (IMP) [36]. Purine metabolism plays an important role in microorganisms, recycling carbon and nitrogen compounds and regulating metabolism [37]. Several previous reports have described the importance of nucleotide biosynthesis by bacteria triggering infections. For instance, certain auxotroph mutants of *Salmonella*, *S. aureus,* or *S. pneumoniae* were avirulent in murine infection models [38, 39]. Besides, other studies demonstrate that limiting amounts of nucleotide bases in human serum, force pathogens to rely on *de novo* nucleotide biosynthesis [40].

Alc, which encodes an allantoicase, is involved in (S)-allantoin degradation and catalyzes the conversion of allantoate to (S)-ureidoglycolate and urea. Previous studies have found that nitrogen controls several pathways involved in secondary metabolism [41]. In general, secondary metabolites are produced by the cells as a response to environmental cues. For example, a notable decrease in antibiotic production was observed in *S. coelicolor* M145 as a response to the excess of intracellular ammonium generated during degradation of allantoin when the microorganism was grown in allantoin as the sole nitrogen source [42].

Regarding proteins involved in amino acids biosynthesis, LeuD and ProA are present in higher amounts in the presence of blue light (FC_LeuD_= 1.62, FC_ProA_= 1.50). LeuD catalyzes the isomerization between 2-isopropylmalate and 3-isopropylmalate, and is part of L-leucine biosynthesis [43].

ProA (gamma-glutamyl phosphate reductase protein) participates in step 2 of the subpathway that synthesizes L-glutamate-5-semialdehyde from L-glutamate and is involved in L-proline biosynthesis [44]. There are shreds of evidence that proline metabolism has complex roles in a variety of biological processes, including cell signaling, stress protection, and energy production [45]. Proline can also contribute to the pathogenesis of various disease-causing organisms. Besides, some organisms use proline directly for the biosynthesis of secondary metabolites with antibacterial or antifungal properties [46].

An interesting protein that is over-represented in the light state is SurA, a chaperone involved in the correct folding and assembly of outer membrane proteins (OMPs), and may act both in early periplasmic as well as in late outer membrane-associated steps of protein maturation [47]. A great variety of outer membrane proteins are porins and autotransporters that facilitate transport and other essential functions, and act as virulence factors [48]. In *P. aeruginosa,* a significantly lower amount of many porins were detected in the outer membrane (OM) of the conditional *surA* mutant, including members of the OprD family (OpdO, OpdN, OpdP, and OprD) [49]. Also, it was found that siderophore receptors were absent or less abundant in the OM upon depletion of SurA [49]. Related to this point, in the present work a TonB-dependent receptor and a biopolymer transporter ExbD, which are part of a complex that energizes specific high-affinity receptors involved in regulating iron uptake, were found in a greater amount under blue light (FC_TonB_= 27.35, FC_ExbD_= 9.27). Consistently, a more robust growth under iron-deprived conditions, i.e., in the presence of the iron chelator 2,2′-dipyridyl (DIP), was observed in *A. nosocomialis* under blue light at 37 °C while it was severely affected in the dark [9]. Therefore, the higher abundance of SurA, TonB-dependent receptor, and ExbD under blue light could contribute to the enhanced iron acquisition observed in this condition, and this is an important feature since iron is a pathogenicity determinant essential for the success of bacterial infections [50].

Other proteins such as queuine tRNA-ribosyltransferase (Tgt) and ribosomal RNA large subunit methyltransferase K/L (RlmL) showed an increase in abundance under blue light. Tgt catalyzes the exchange of a guanine 34 with the queuine precursor 7-aminomethyl-7-deazaguanine (PreQ1) in specific tRNAs containing anticodones G(guanine)-U(uracil)-N (tRNA-Asp, -Asn, -His and -Tyr), where N is one of the four canonical nucleotides [51]. It was shown that a functional Tgt is required for efficient pathogenicity of *Shigella* bacteria [52]. A null-mutation in the *tgt* gene strongly reduces translation of *virF*-mRNA, a transcriptional activator required for the expression of a large number of *Shigella* pathogenicity genes [53]. On the other hand, RlmL K/L specifically methylates the guanine in position 2445 and the guanine in position 2069 (m7G2069) of 23S rRNA. The rRNA methyltransferases (rRNA MTases), especially those acting on 23S rRNA, are associated with development of antibiotic resistance in a wide variety of bacteria, and some of them are recurrent human pathogens [54]. Inactivation of 23S rRNA MTases function has been shown to affect negatively translation and cell physiology [55].

Finally, a glutathione S-transferase was over-represented in the light condition. The glutathione S-transferases (GSTs) are a family of proteins that conjugate glutathione to the sulfur atom of cysteine in various compounds [56]. GST can bind to a variety of hydrophobic compounds endogenous and xenobiotic alkylating agents with high affinity, including carcinogens, therapeutic drugs, environmental toxins and products of oxidative stress, protecting cells from oxidative damage [57].

Taken together, these observations suggest that light could play a main role in the control of *A. nosocomialis* physiology at 37 °C, particularly modulating pathogenesis and allowing cells to respond and adapt to environmental signals.

### Proteins with increased abundance in the dark compared to light

Among the proteins with higher abundance in the dark is present the phenylacetate-CoA oxygenase, PaaK subunit. Phenylacetate-CoA oxygenase is comprised of a complex composed of 5 proteins responsible for the hydroxylation of phenylacetate-CoA (PA-CoA), which is the second catabolic step in phenylacetic acid (PAA) degradation [58]. Interestingly, we have shown that *A. nosocomialis* RUH2624 growth is stimulated in the dark when PAA is present as the sole carbon source at 37 °C [9], supporting the notion that the PAA catabolic pathway is modulated by light. Modulation of the PAA catabolic pathway has been shown to influence *A. baumannii*’s pathogenesis. In fact, inhibition of this pathway resulted in increased neutrophil migration to the site of infection and bacterial clearance [58] [59].

Another protein that appears over-represented in the dark is the nonhydrolyzing UDP-N-acetylglucosamine 2-epimerase (FC= 1.7). This enzyme catalyzes the reversible interconversion of UDP-N-acetylglucosamine (UDP-GlcNAc) to UDP-N-acetylmannosamine (UDP-ManNAc) [60], being this compound an intermediate in the biosynthesis of a variety of bacterial capsular polysaccharides (CPSs) [61]. In several pathogenic strains, such as *B. anthracis*, *N. meningitides* and *S. aureus*, CPSs are important virulence factors that protects bacteria from the immune system of a host and harsh environmental conditions [62–64]. In this sense, the UDP-N-acetylglucosamine 2- epimerase could contribute to cell-surface polysaccharide synthesis in *A. nosocomialis,* protecting the cells in the absence of light. The probable potassium transport Kup protein, found in a greater quantity in dark, could also favor the adaptation to rapidly changing external conditions.

## Conclusion

In this work, we show that light regulates motility in *A. nosocomialis* in a wide range of temperatures, which go from environmental to temperatures found in warm-blooded hosts (23 to 37 °C). This temperature dependence is different from that observed for regulation of motility by light in *A. baumannii* ATCC 17978, which was shown to occur only in the low to environmental temperature range (18 to 24 °C). We hypothesized that this could be due to an unequal endowment of *blsA* homologs, whose intrinsic characteristics may allow them to function in other temperature ranges. In this work, we show that the three BLUF domain-containing genes present in the *A. nosocomialis* RUH2624 genome [8], encode active photoreceptors. But only two of them, AnBLUF65 and AnBLUF46 are stable proteins when produced *in vitro* in the temperature range analyzed: (15-37 °C) and (15-32 °C), respectively. The fact that AnBLUF65 is the only of the three BLUF domain-containing proteins that is expressed *in vivo* along with the photo-regulatory temperature range, strongly suggests that it participates in the modulation of motility by light. Spectroscopic characterizations of AnBLUF65 and AnBLUF46 *in vitro*, indicate that AnBLUF65 is more efficient to form the light-adapted state than AnBLUF46 at 24 °C. And although AnBLUF65 efficiency is negatively affected by temperature increments, the protein remains active at 37 °C, accordingly with being the only blue light photoreceptor expressed in *A. nosocomialis*. 3D models of these proteins were presented and discussed. The relative solvent accessibility to the cofactor pocket derived from flavin emission fluorescence correlates well with the tertiary structure modeled for both proteins and both states, dark and light-adapted. We have additionally characterized intramolecular interactions to profile the underlying phenomena upon formation of the light-adapted state. The presence of aromatic clusters networks on either side of the β-sheet has been described. We propose that the rupture of an Asp-X interaction via H-bonding observed upon illumination in the loop between the Tyr-Tyr and the first set of Phe-Phe motifs, allows the aromatic tetrads in the variable domain to displace and to adopt the light-adapted state of this part in AnBLUF65 and AnBLUF46 (Fig 10). Thus, the calculated strengthening of the π-π tetrads in AnBLUF65 and the displacements of these residues between the dark and the light-adapted forms in BlsA as observed by Chitrakar et al 2020, reinforce the idea that intramolecular signaling events involve, at least in part, hydrophobic/aromatic interactions. The extent to which these networks are present in other members of the genus *Acinetobacter*, and other microorganisms remain to be further explored.

**Fig 10:**
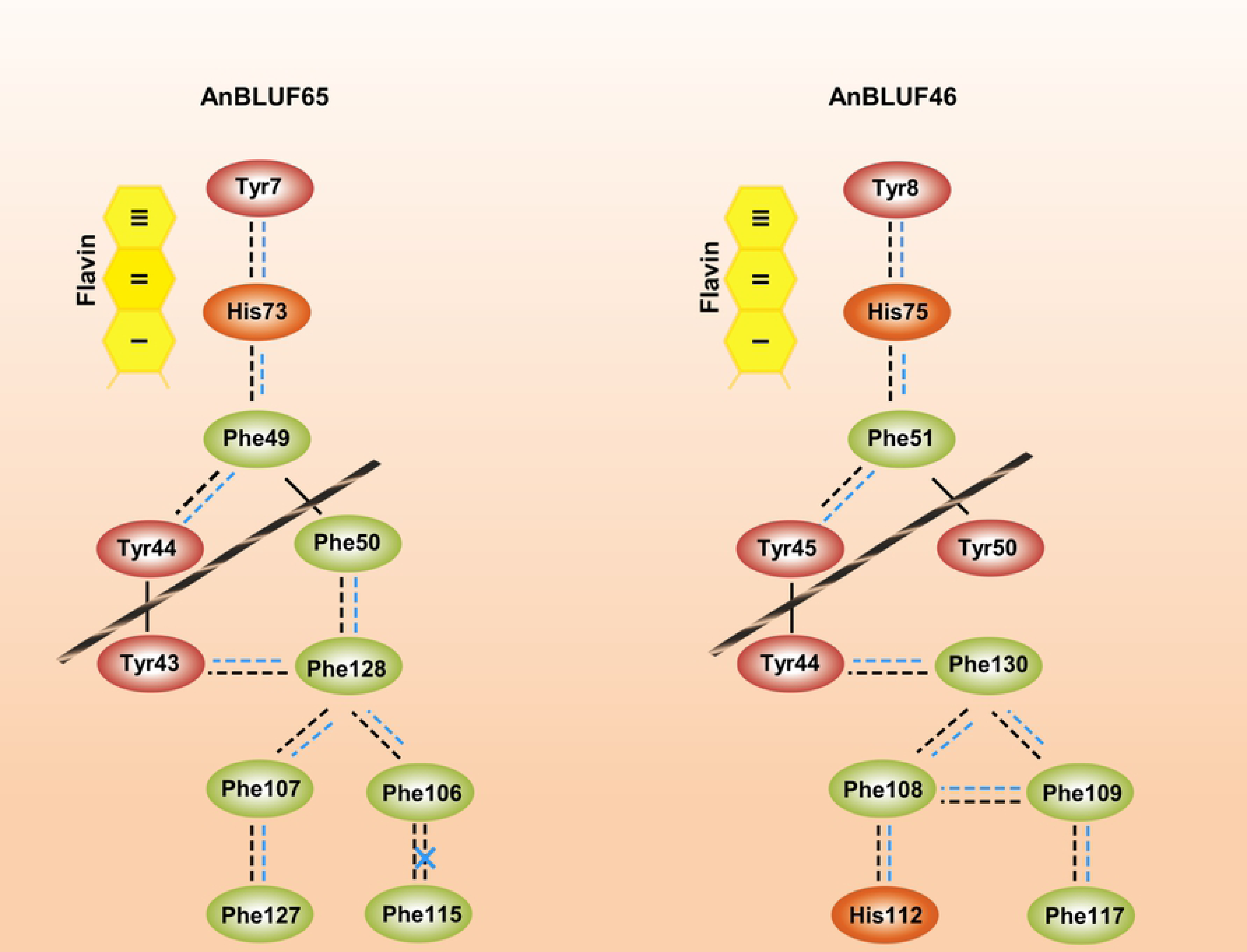
Intra-protein aromatic networks in BLUFs from *A. nosocomialis*. Black ribbons represent β sheet dividing protein in two areas: Flavin/N-terminal area and variable domain. Solid lines correspond to main chain covalent bonds. Dashed lines represent relative distances between aromatic centroids for dark (black) and blue light (blue).

Overall, in this work, we have characterized different BLUF photoreceptors and show that they are operative at different temperature ranges. Despite diverse BLUF-coding genes are encoded in *A. nosocomialis*, not all are physiologically functional. It is worth mentioning that AnBLUF65 is the only BLUF-protein found to be expressed in *A. nosocomialis*, and the possibility raises that the other BLUF-photoreceptors are expressed under different conditions, such as growth media, temperatures, presence of a host, etc. The presence of AnBLUF85 only in two different *A. nosocomialis* strains and discretionally in other species of the *Acinetobacter* genus is striking, in addition to the fact that this photoreceptor is not stable and not expressed in its host. The overall information suggests that this gene might be cryptic; despite it does not seem to be subjected to genetic drift as its sequence is conserved in different species, and questions arise regarding its evolutionary origin and functionality.

## Acknowledgment

This work was supported by grants from the Agencia Nacional de Promoción Científica y Tecnológica (PICT 2018- 00793) to MAM, the Consejo Nacional de Investigaciones Científicas y Tecnológicas (PIP 2017-0493) to IA and ASaCTeI (Ministerio de Ciencia, Tecnología e Innovación Productiva de la Provincia de Santa Fe) IO-021-18 to MAM. We would like to thank Dr. Javier M. González for his critical reading of the manuscript and suggestions. Also, Lic. Alba Loto for her assistance with protein overexpression and purification. IA, CDB and MAM are career investigators of CONICET. BPM and MRT are fellows from the same institution. LV is a Universidad Nacional de Santiago del Estero investigator.

## Supporting information

**S1 Fig.**
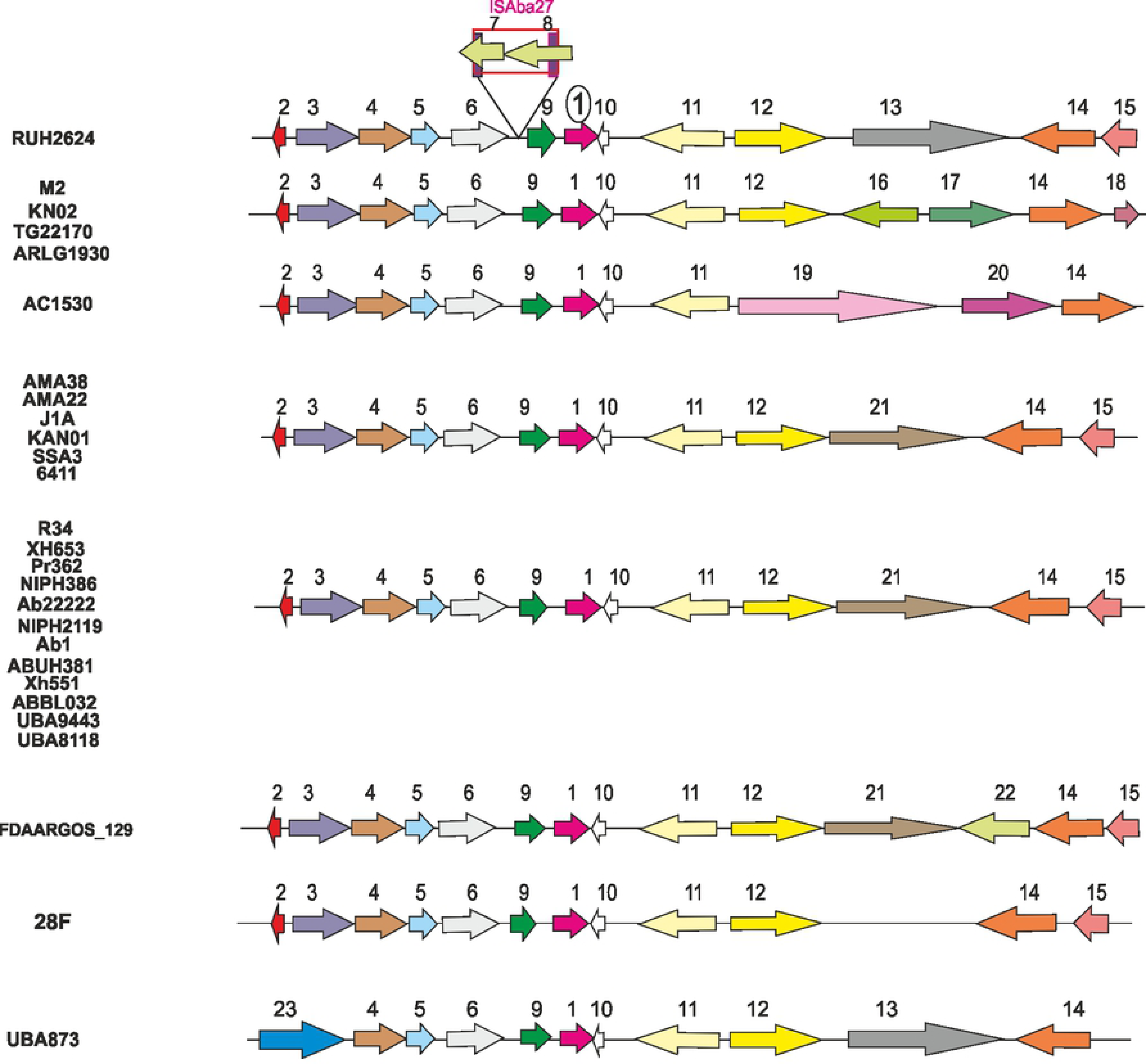
Genomic context of AnBLUF65 homologs present in different strains of *A. nosocomialis*. (A). Coding-sequences (CDSs) are located in their corresponding frame. AnBLUF65 is indicated as pink arrow. Different colors indicate different functions. Gene annotations are indicated as numbers above the schemes, with the following codes: 1- AnBLUF65, 2- DUF2171, 3- Acyl-CoA-dehydrogenase, 4- GlcNAc-PI-de-N-acetylase, 5- methyltransferase domain, 6- glycosyltransferase, 7- DDE endonuclease domain, putative transposase, 8- helix-turn-helix fo DDE superfamily endonuclease, 9- BOF- class 2b aminoacyl-tRNA synthetases- NirD/YgiW/Y damily stress tolerance protein, 10- HP, 11- poly (R)-hydroxyalkanoic acid synthase, 12- sodium/glutamate symporter, 13- proton antiporter-2 (CPA2) family, 14- NDAB- Rossmann Superfamily- Oxidoreductase, 15- DoxX-like family, 16- LysR family transcriptional regulator, 17- AKR15A family of aldo-keto reductase, 18- pyrabactin resistance 1 (PYR1) receptor, 19- type I secretion target GGXGXDXXX repeat protein, 20- Paax domain, 21- glutathione-regulated potassium-efflux system protein KefC, 22- IS3 family transposase, 23- L- asparagine transporter.

**S2 Fig.**
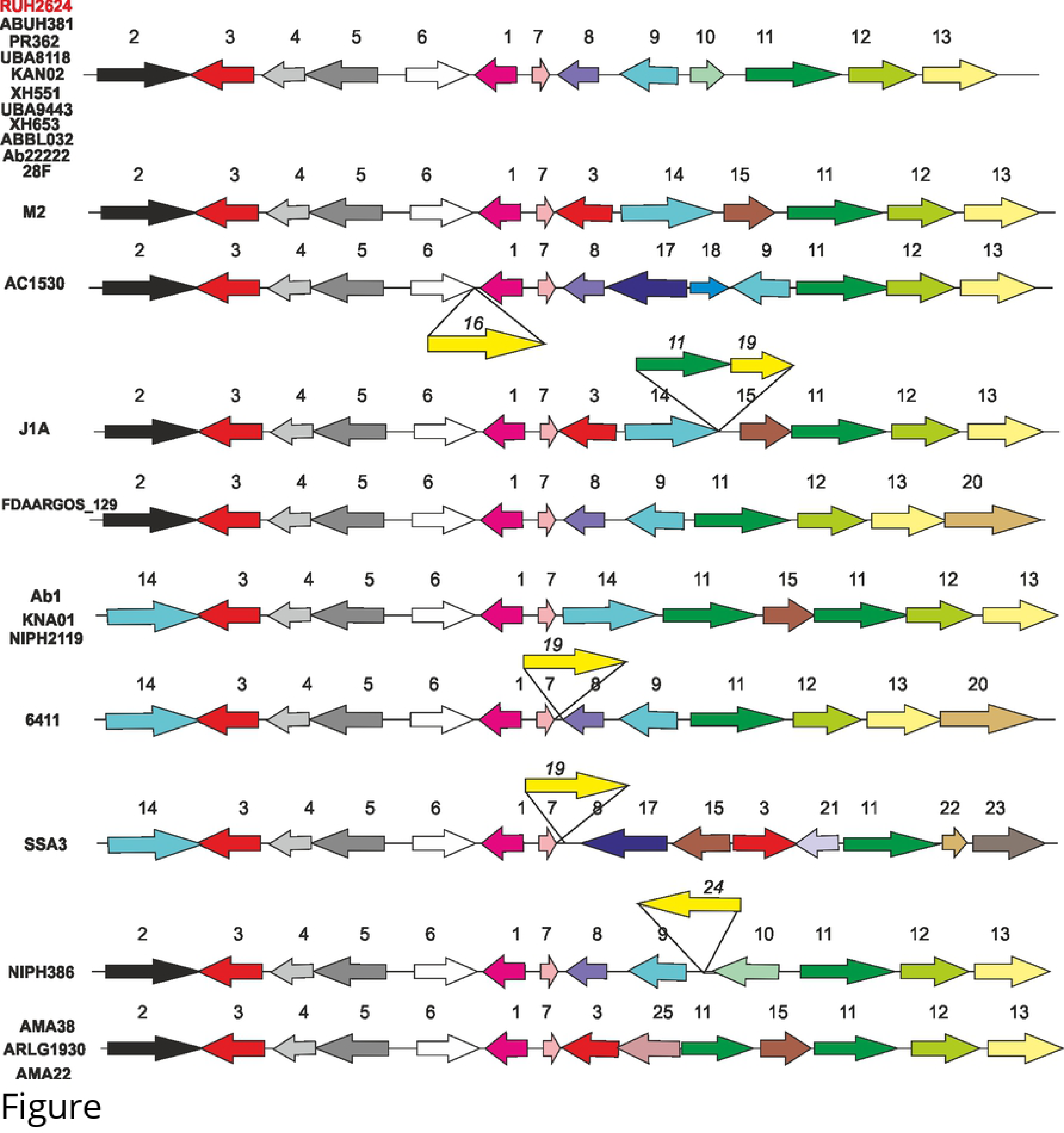
Genomic context of AnBLUF46 homologs present in different strains of *A. nosocomialis*. (A). Coding-sequences (CDSs) are located in their corresponding frame. AnBLUF46 is indicated as pink arrow. Different colors indicate different functions. Gene annotations are indicated as numbers above the schemes, with the following codes: 1- AnBLUF46, 2- aromatic acid:H^+^ symporter (AAHS), 3- AraC family transcriptional regulator RmlC-cupin protein, 4- 3- hydroxyburyrate dehydrogenase, 5- H^+^/gluconate symporter, 6- MBL fold metallo-hydrolase, 7- HP, 8- DUF2846- putative conjugal domain, 9- DUF3298- putative heat shock protein, 10- HP, 11- LysR family transcriptional regulator, 12- succinyl-CoA:3- ketoacid-coenzyme A transferase subunit A, 13- succinyl-CoA:3-ketoacid-coenzyme A transferase subunit B, 14- multidrug efflux MFS transporter, 15- pimeloyl-ACP methyl ester carboxylesterase, 16- IS5 transposase, 17- TetR/AcrR family transcriptional regulator, 18- DoxX family protein, 19- IS transposase, 20- anion permease ArsB/NhaD, 21- 3-oxoacyl-ACP reductase FabG, 22- nuclear transport factor 2 family protein, 23- fermentarion-respiration switch protein FrsA, 24- DDE transposase domain, 25- Bcr/CflA family drug resistance efflux transporter.

**S3 Fig.**
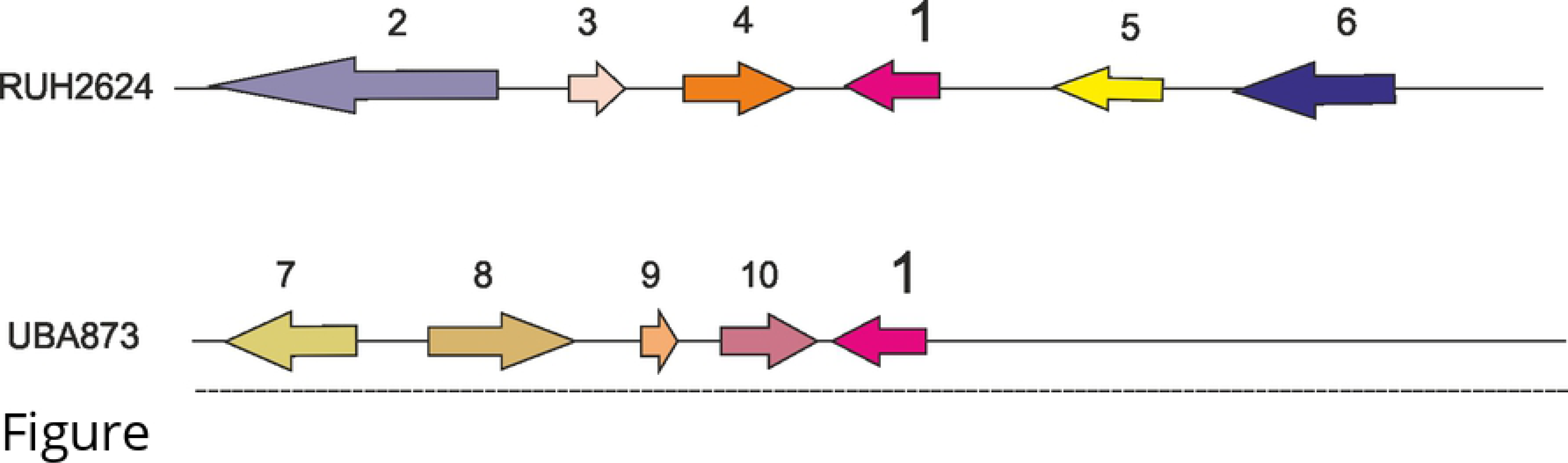
Genomic context of AnBLUF85 homologs present in different strains of *A. nosocomialis*. (A). Coding-sequences (CDSs) are located in their corresponding frame. AnBLUF85 is indicated as pink arrow. Different colors indicate different functions. Gene annotations are indicated as numbers above the schemes, with the following codes: 1- AnBLUF85, 2- relaxase-MobA/MobL family protein, 3- FAM199X, 4-HP, 5- N-acetiltransferasa, 6- conjugal transfer pilus assembly protein TraB, 7-RepB initiator replication protein, 8- potasium transporter, 9-mRNA-degrading endonuclease RelE, 10- chromate transport protein ChrA.

**S4 Fig:**
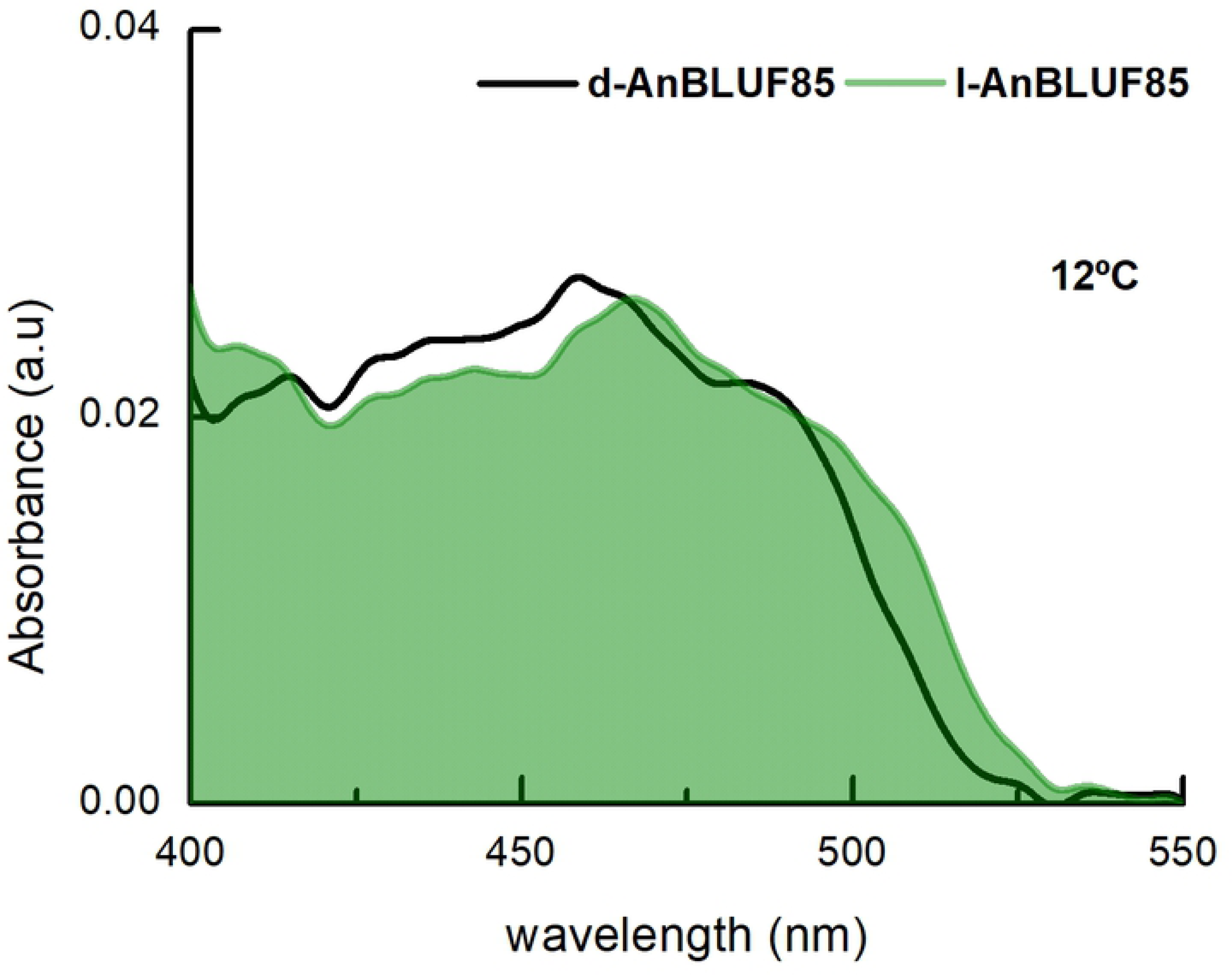
Visible absorption spectra changes of AnBLUF85 photoreceptor by blue light illumination, in working buffer solution at 12 °C. dAnBLUF85 (black line) and lAnBLUF 85 (pale green)

